# Retinoblastoma protein activity revealed by CRISPRi study of divergent Rbf1 and Rbf2 paralogs

**DOI:** 10.1101/2023.05.19.541454

**Authors:** Ana-Maria Raicu, Patricia Castanheira, David N. Arnosti

## Abstract

Retinoblastoma tumor suppressor proteins regulate the key transition from G1 to S phase of the cell cycle. The mammalian Rb family comprises Rb, p107, and p130, with overlapping and unique roles in gene regulation. Drosophila experienced an independent gene duplication event, leading to the Rbf1 and Rbf2 paralogs. To uncover the significance of paralogy in the Rb family, we used CRISPRi. We engineered dCas9 fusions to Rbf1 and Rbf2, and deployed them to gene promoters in developing Drosophila tissue to study their relative impacts on gene expression. On some genes, both Rbf1 and Rbf2 mediate potent repression, in a highly distance-dependent manner. In other cases, the two proteins have different effects on phenotype and gene expression, indicating different functional potential. In a direct comparison of Rb activity on endogenous genes and transiently transfected reporters, we found that only qualitative, but not key quantitative aspects of repression were conserved, indicating that the native chromatin environment generates context-specific effects of Rb activity. Our study uncovers the complexity of Rb-mediated transcriptional regulation in a living organism, which is clearly impacted by the different promoter landscapes and the evolution of the Rb proteins themselves.

## INTRODUCTION

The Retinoblastoma (Rb) tumor suppressor protein is an ancient and highly conserved eukaryotic transcriptional corepressor. Rb is a member of a family of pocket proteins that includes the p107 and p130 paralogs. These proteins share a conserved central pocket domain, through which they bind to E2F transcription factors that are found on gene promoters (Zhang and Dean, 2001). Rb binding to the E2F transactivation domain allows for transient and reversible inhibition of E2F activity, which leads to the nearby gene’s repression. At a secondary site in the pocket domain, Rb proteins recruit cofactors such as histone modifiers and chromatin remodelers, leading to chromatin modifications and subsequent repression of target genes (Zhang and Dean, 2001).

Canonical Rb targets are cell cycle genes; in particular, Rb regulates genes involved in the progression from G1 to S phase of the cell cycle. In G1, Rb represses these cell cycle genes through E2F binding; phosphorylation of Rb relieves the repression, allowing for progression into S phase. Rb proteins also regulate the expression of genes involved in diverse processes including DNA repair, transcription, apoptosis, polarity, signaling, and metabolism (Reviewed in Chau and Wang, 2003, Dick and Rubin, 2013, Nicolay and Dyson, 2013; Payankaulam *et al*. 2016).

Rb was the first tumor suppressor protein to be identified in humans, as loss of heterozygosity of the *RB1* gene leads to pediatric retinoblastoma (Knudson, 1971). Mutations in the Rb pathway (including E2F, cyclins, and Cdk proteins) have been identified in most human cancers, implicating this pathway in both the initiation and the progression of cancer. Although all Rb proteins function in similar ways as transcriptional corepressors, Rb is more often found to be mutated in human cancer than p107 and p130, and is considered the predominant tumor suppressor (Weinberg, 1992). Rb plays a leading role in gene regulatory processes, while p107 and p130 have some tissue-specific or promoter-specific roles, and do not substitute for the full spectrum of Rb activity in Rb null tissues.

All vertebrates encode at least three Rb proteins, with some expressing up to six (Liban *et al*. 2017). Interestingly, like the vertebrate lineage, the Drosophila genus experienced an independent Rb gene duplication event; this dates back ∼50 million years and is the only Rb duplication recorded within Arthropoda. The selective advantage for multiple RB genes in both vertebrates and in the Drosophila lineage is not well-understood. *Drosophila melanogaster* is an excellent system for studies aimed at uncovering the evolutionary significance of Rb paralogy, and determining how gene regulatory tasks are apportioned between Rb family members.

Initial studies of the Drosophila Rbf1 and Rbf2 paralogs identified major differences between the paralogs; Rbf1 was deemed the predominant corepressor, with Rbf2 a weaker version of Rbf1. *rbf1* null flies are larval lethal, but *rbf2* null flies are viable, exhibiting fertility and lifespan defects (Du and Dyson, 1999; Steveaux *et al*. 2002; Steveaux *et al*. 2005; Mouawad *et al*. 2019). Additionally, in the adult, Rbf1 is expressed widely, while Rbf2 is localized mostly to the ovary (Stevaux *et al*. 2002, Keller *et al*. 2005). Indeed, we and others have shown that when assayed on specific cell cycle promoters, Rbf1 is a much more potent repressor than Rbf2 (Steveaux *et al*. 2002, Mouawad *et al*. 2019). Additionally, suppression of *rbf2* by RNAi in S2 cells does not lead to misregulation of many genes, but *rbf1* depletion does lead to upregulation of many genes, including cell cycle genes (Dimova *et al*. 2003). Despite its weaker activity in these contexts, we found that Rbf2 binds to twice as many genomic targets as Rbf1 in embryos, with an enrichment of mitochondrial protein and ribosomal protein genes being specifically bound and regulated by the Rbf2 paralog (Acharya *et al*. 2012, Wei *et al*. 2015, Mouawad *et al*. 2019). Additionally, we recently showed that Rbf2 is a stronger repressor than Rbf1 on specific promoters, such as *tko* and *cycB* (Wei *et al*. 2015; Mouawad *et al*. 2020).

Although Rb proteins have been found to interact with sites distal to gene promoters, binding is often close to transcriptional start sites (TSS). Genome-wide ChIP studies performed on Drosophila embryos identified preferential binding within 500 bp of a gene’s TSS, with a peak at -200 bp (Acharya *et al*. 2012; Wei *et al*. 2015). ChIP-seq in Drosophila third instar larvae and in human senescent cells confirmed the promoter-proximal preference of Rb proteins (Korenjak *et al*. 2012; Chicas *et al*. 2010). The significance of this binding preference remains an outstanding question. Rb proteins may directly target the basal machinery as short-range repressors, or they may be constrained by preferential binding to the E2F transcription factors, which themselves are promoter-associated. The recent mapping of Rb proteins to enhancers and insulators suggests additional complexity, at least in mammals (Sanidas *et al*. 2022).

Whether Rb functions as a short-range or long-range repressor on endogenous loci has not been definitively answered. The impact of short-versus long-range repression can have profound consequences for gene expression: long-range repressors can impact multiple distal regulatory elements for long-distance silencing whereas short-range repressors act locally to provide precise, tunable regulatory effects (Gray and Levine 1996; Barolo and Levine, 1997; Li and Arnosti, 2011). Initial studies tethering Rb to the GAL4 DNA binding domain allowed for testing the ability of Rb proteins to function from proximal or distal positions. In transfection assays, GAL4-Rb can repress an SV40 enhancer from both proximal and distal positions up to 2 kb away, and from both upstream and downstream sites relative to the TSS, suggesting a possible long-range repression mechanism (Weintraub *et al*. 1995; Bremner *et al*. 1995; Adnane *et al*. 1995; Luo *et al*. 1998). More recently, the Stark lab used GAL4-Rb fusions along with STARR-seq in Drosophila S2 cells to compare the impact of Rbf1 and Rbf2 paralogs. These studies revealed that thousands of enhancers can be repressed by the Drosophila paralogs from a position adjacent to the tested regulatory element, but distal to the TSS, indicating that TSS proximity is not essential for repression and that Rb also has repressive effects on enhancers (Jacobs *et al*. 2022, bioRxiv). Yet, such transiently transfected reporters may not fully recapitulate Rb biology on endogenous loci.

Instead of relying on global perturbations or genome-wide binding assays, we turned to CRISPRi for comparative analysis of Rbf1 and Rbf2, thus providing a method to answer questions about intrinsic repression activity of the paralogs at diverse genomic loci. Using gene-specific guide RNAs, we targeted dCas9-Rb chimeras to a variety of promoters in a tissue-specific manner. We find that Rbf1 and Rbf2 have diverse effects on promoters; in some cases, the two paralogs have similar effects, while on other promoters one paralog is more potent than the other. The CRPSPRi system also permitted a structure-function analysis of Rbf1, revealing the role of the pocket domain and instability element for promoter targeting, but not repression activity. Overall, we find that Rb proteins are not promiscuous, dominant corepressors; rather, many times they work as soft repressors, modestly dialing down gene expression rather than functioning as an on/off switch. Additionally, they are highly responsive to the specific promoter landscape in which they act. Our identification of context-specific roles of Rb paralogs *in vivo* points to differences in intrinsic repressive activity, shaped by evolutionary changes in Rb paralogs over time.

## RESULTS

### Deploying dCas9-Rb chimeras in Drosophila

To investigate comparative Rb paralog activity on endogenous genes in Drosophila, we created FLAG-epitope-tagged dCas9-Rbf1 and dCas9-Rbf2 chimeras, and FLAG-tagged dCas9 alone as a negative control (**Figure 1**). We also employed the dCas9-VPR chimera, which functions as a strong activator in Drosophila (Ewen-Camben *et al*. 2017). dCas9-Rb chimeric proteins have not been tested before in any system, necessitating a proof-of-principle approach in which we could quickly screen dCas9-Rb impacts *in vivo*. To test the impact of dCas9-Rb recruitment to gene promoters, we used the GAL4/UAS system for tissue-specific expression of the chimeras in the developing fly wing. Specifically, we used the *nubbin* driver, which is expressed predominantly in the L3 wing disc, to screen for transcriptional impacts in larvae and phenotypic impacts in the adult wing. To this end, we created homozygous transgenic fly lines expressing *nub*-GAL4 and one of the UAS:dCas9-Rb effectors. The *nub*-GAL4>UAS:dCas9-Rb flies were crossed to homozygous flies expressing two tandem gRNAs for a gene’s promoter, to create flies expressing a single copy of each of the three transgenes (**Figure 1B**; see Materials & Methods). We confirmed genotypes of flies by PCR and tested mRNA expression level of the transgenes in the wing disc using RT-qPCR (**Figure 1C; S1B, C**). The dCas9-Rb effectors are expressed at similar levels of mRNA and also at similar protein levels *in vivo* (**Figure S1**). Crossing *nub*-GAL4>UAS:dCas9-Rb flies to a non-targeting gRNA control fly line (QUAS; Ewen-Camben *et al*. 2017), we observed phenotypically wild type (WT) adult wings, indicating that expression of a single copy of each of the transgenes does not impact normal wing development (**Figure S2F**).

**Figure 1.**
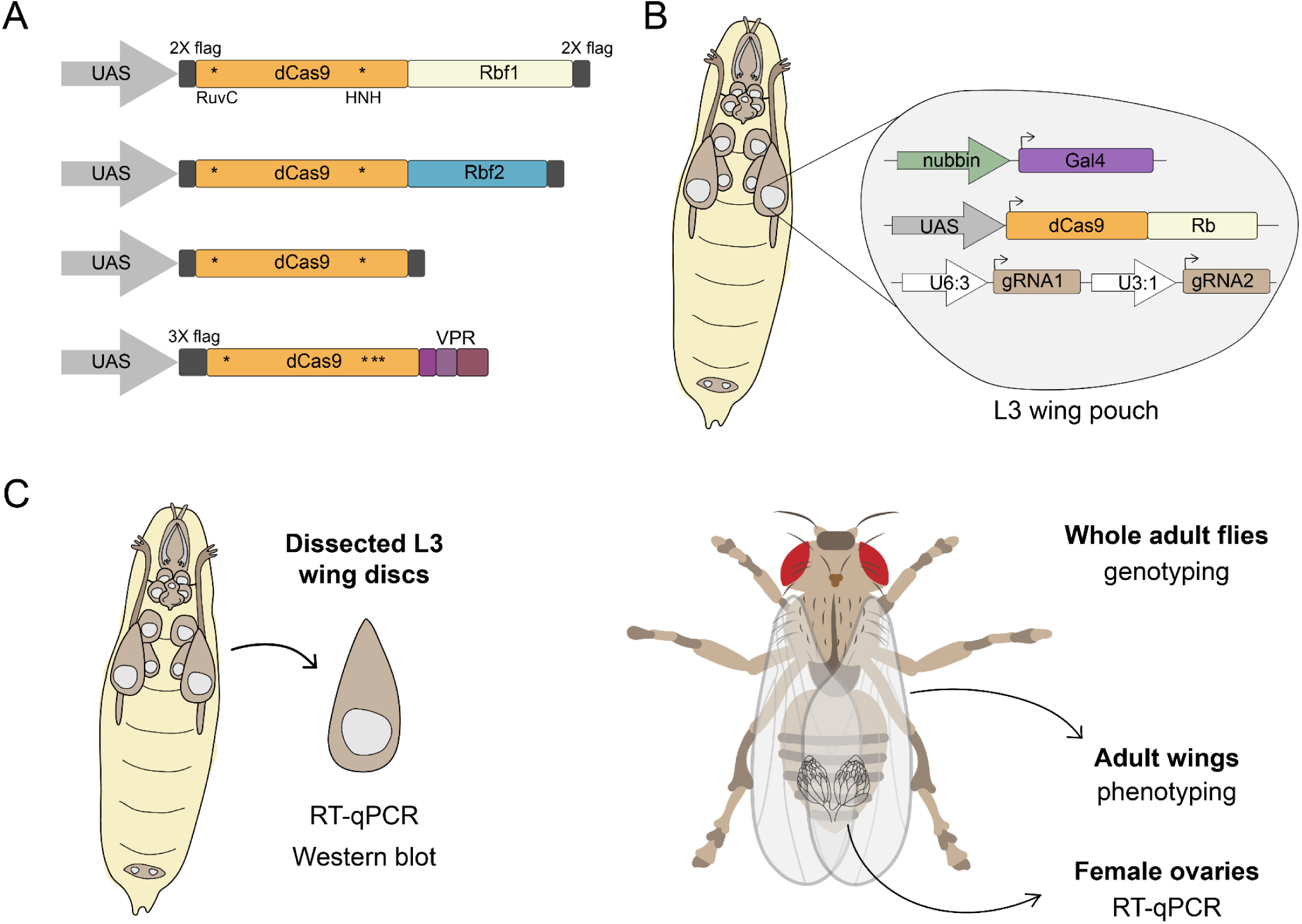
Creation of an *in vivo* system for targeting Rb paralogs to gene promoters using CRISPRi. **A)** The fly Rbf1 and Rbf2 FLAG-tagged coding sequences were fused to the C-terminus of the *S. pyogenes* nuclease dead Cas9 (dCas9; D10A mutation in RuvC catalytic domain and H840A mutation in HNH catalytic domain), and placed under UAS expression. FLAG-tagged dCas9 was created as a negative control. dCas9-VPR was obtained as a fly line; VPR is a tripartite activator that was previously characterized as a Drosophila transcriptional activator (Ewen-Campen *et al*. 2017). Asterisks indicate mutation sites in dCas9. **B)** *Drosophila melanogaster* expressing three transgenes were generated for tissue-specific expression of dCas9-Rb effectors using GAL4-UAS. Flies express dCas9-Rb chimeras in the *nubbin* expression pattern (begins in L2 and then localized to the wing pouch of L3 wing discs), with ubiquitous expression of two tandem gRNAs designed to target a single gene’s promoter. Flies used in experiments express one copy of each of the three transgenes. **C)** Schematic of experimental procedures performed with different developmental stages. L3 wing discs were used for RT-qPCR analysis of gene expression changes and measuring mRNA and protein expression levels of the dCas9 effectors. Whole adult flies were used for genotyping, adult wings for phenotyping, and female ovaries for RT-qPCR analysis of gene expression changes.

### Gene-specific effects of Rbf1 and Rbf2 recruitment by dCas9

Studies of Rb paralogs indicate that Rbf1 and Rbf2 are not fully redundant, and may have promoter-specific effects (Wei *et al*. 2015; Mouawad *et al*. 2019). For instance, Rbf1 is found on ∼2000 gene promoters in the embryo, while Rbf2 is found on ∼4000 promoters. Additionally, Rbf1 associates with both dE2F1 and dE2F2 while Rbf2 only interacts with dE2F2 (Stevaux *et al*. 2002). The molecular basis for preferential recruiting of one paralog versus the other is unresolved. To test the impact of each Rb paralog on endogenous promoters, we recruited them to diverse loci as described above (**Figure 1B**). We tested promoters of 28 genes, which are either known targets of one or both Rb paralogs, or genes with no known Rb regulation based on ChIP-seq and RNA-seq data (**Supplementary Table 1**; Wei *et al*. 2015; Mouawad *et al*. 2019). The selected genes span a variety of functions, including cell cycle regulation, DNA replication, signaling pathways, and ribosomal protein genes. Overall, wing phenotypes differed based on the targeted gene, with some promoters only responsive to the repressors, or the activator, or both. In some instances, the phenotype was fully penetrant, while in others, only a subset of collected wings displayed the observed phenotype (**Supplementary Table 1**).

Here, we detail the effects on eight of these genes, where stronger morphological phenotypes were observed: *E2F2, Mpp6* (cell cycle), *InR*, *wg*, *dpp* (signaling pathways), *Acf* (histone remodeler), *mcm6* (DNA replication), and *Pex2* (peroxisome protein). Adult wing phenotypes ranged from mild (missing anterior or posterior crossvein, ACV and PCV) to moderate (supernumerary bristles, ectopic venation) to severe (severe morphological defect; **Figure 2, S2**).

**Figure 2.**
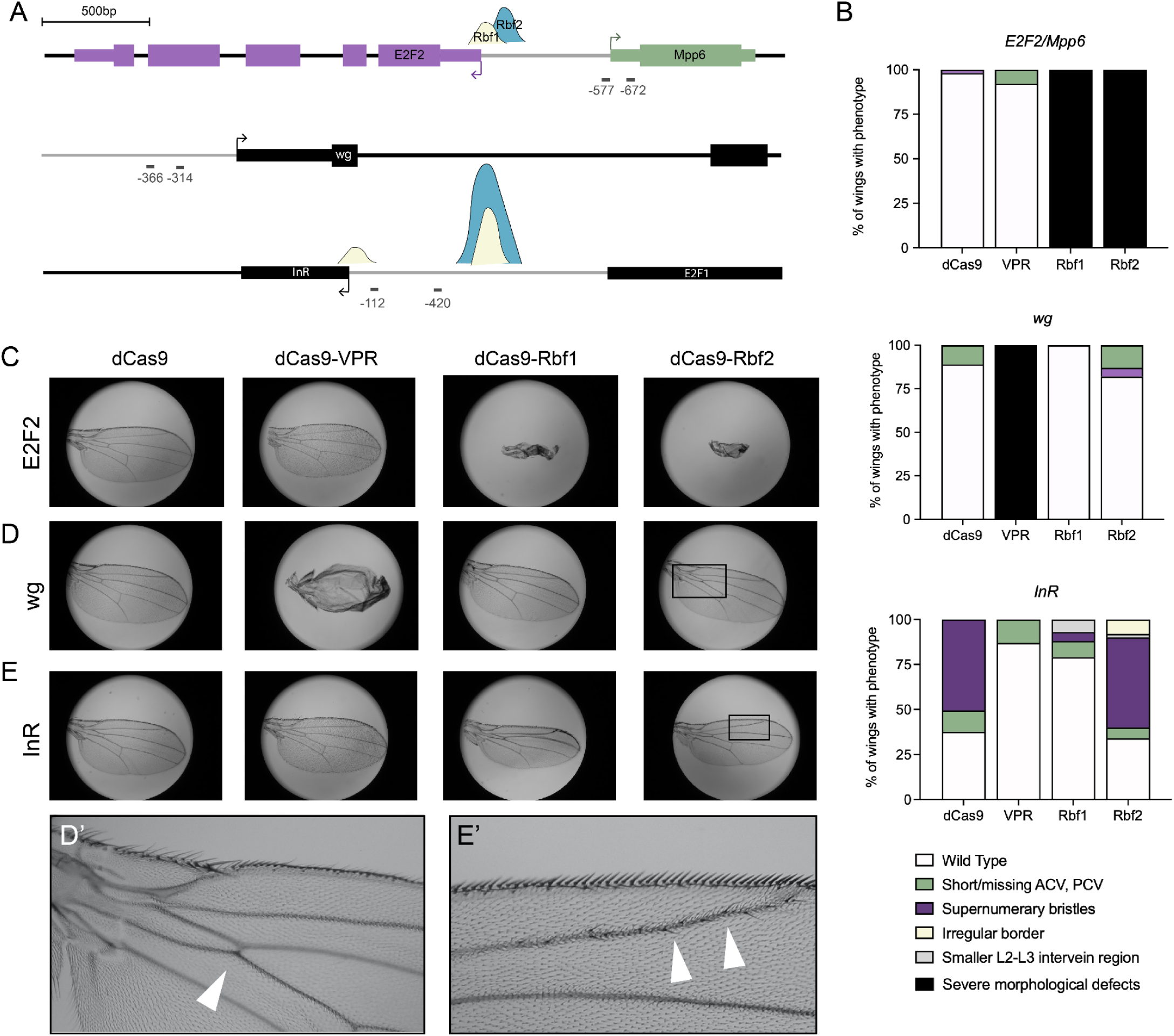
Rbf1 and Rbf2 targeting to gene promoters in the wing leads to unique phenotypic effects based on target gene. **A)** Diagram of gRNA binding sites on the promoters of *E2F2*, *wg,* and *InR*. Arrows indicate the TSS. gRNAs targeting *E2F2* overlap with the nearby gene, *Mpp6*. Thick black bars are exons, thinner bars are Untranslated Regions (UTR), and black horizontal lines are introns. Gray horizontal lines indicate intergenic regions, and short lines below the genes are the locations of the gRNAs, with approximate distance from the indicated TSS below. These gRNAs were obtained from Harvard TRiP (Zirin *et al*. 2020). Peaks indicate Rb binding sites from embryo exo-ChIP (Wei *et al*. 2015). **B)** Adult wings from ∼50 flies (n=100) in which dCas9 effectors were targeted to the *E2F2, wg,* and *InR* genes were analyzed for phenotypic effect after targeting. Legend below indicates the observed phenotypes. Values are indicated as a proportion of the total number of wings. **C)** Representative images from targeting the *E2F2* promoter. Rbf1 and Rbf2 led to severe morphological defects with 100% penetrance. In contrast, dCas9 alone and dCas9-VPR targeting produced WT wings. **D)** Representative images from targeting the *wg* locus. The VPR activator led to severe morphological defects with 100% penetrance, while Rbf2 had mild effects and Rbf1 had no impact on wing development. **D’** Inset indicates the “short ACV” phenotype seen with Rbf2. **E)** Representative images from targeting the *InR* locus. All effectors caused mild effects, including dCas9 alone. Here, effects from dCas9 and dCas9-Rbf2 are indistinguishable, while Rbf1 targeting led to a smaller number of wings with a phenotype. **E’** Inset shows the “supernumerary bristles” phenotype seen with many of the effectors.

Targeting dCas9-Rbf1 or -Rbf2 to the *E2F2* promoter leads to a fully penetrant, severe morphological defect (**Figure 2B, C**). This is specific to Rb recruitment, as dCas9 alone does not produce an observable phenotype, nor does the VPR activator. The two gRNAs designed to target *E2F2* actually bind closer to *Mpp6*, which is a divergently transcribed gene from *E2F2*. Therefore, the observed phenotype of Rbf1 and Rbf2 targeting may involve changes in expression of either of these genes, which we explore later. In contrast to these severe Rb-specific effects on *E2F2/Mpp6*, recruitment of the chimeric dCas9-Rb effectors to *wg* does not produce strong, visible phenotypes. Yet, the VPR activator produces severe morphological defects, consistent with previous CRISPRa reports (**Figure 2B, D**; Ewen-Campen *et al*. 2017). As discussed below, the Rb chimeras did produce transcriptional effects.

Targeting the *InR* promoter leads to a diversity of mild to moderate phenotypes, specifically by dCas9 and dCas9-Rbf2 (**Figure 2B, E**). This is an instance where dCas9 alone impacts wing development after binding to the *InR* promoter, perhaps by disrupting the promoter through steric effects. The lack of phenotype from dCas9-VPR may be because the non-specific inhibitory effects produced by dCas9 are ameliorated by the VPR activation domain. Interestingly, the different phenotypic impact by Rbf1 versus Rbf2 suggests that here, Rbf2 may be mediating more potent regulation of *InR* than Rbf1, which we explore later. On the *Acf, Pex2*, *mcm6*, and *dpp* promoters, an assortment of effects are observed (**Figure S2**). In some cases, Rbf1 and Rbf2 are equally penetrant, and in others, they differ in effects. Some of these promoters, such as *dpp*, are more significantly impacted by the VPR activator than by the repressors, while others, such as *mcm6* are equally affected by all the chimeras.

In conclusion, by targeting 28 gene promoters *in vivo*, we show that there are promoter-specific effects, cases where the Rb paralogs can generate the same effect, and instances where they differ in phenotypic impact. We speculate that these context-specific effects may be due to the diversity of *cis*-regulatory elements and *trans*-acting factors on each of these targeted promoters, or because these genes do not play a prominent role in the development of the wing at the targeted stage. Yet, the diversity of phenotypes suggests that the Rb corepressors may work in a context-specific manner; therefore, it is important to dissect the mechanistic differences of these paralogs through assessing transcriptional impact.

### Transcriptional effects of chimeric dCas9-Rb corepressors

To test the transcriptional impact of Rb recruitment on the *E2F2/Mpp6* locus, we collected L3 wing discs and performed RT-qPCR. The tandem gRNAs used above are indicated as gRNA 4 and 5 (**Figure 3A**). Based on previous RNA-seq experiments performed in the embryo and in wing discs, we know that when overexpressed as non-dCas9-fusion proteins, both Rb paralogs misregulate *E2F2* and *Mpp6* to different extents; thus, we expected to observe repression of both genes by the dCas9-fused paralogs (**Supplementary Table 1**). Indeed, both genes were repressed and both Rb effectors showed transcriptional repression activity. *E2F2* was more modestly repressed than *Mpp6* overall, and Rbf1 effects were greater than Rbf2 effects. Both paralogs repressed *Mpp6* >50%, which was greater than the effect of dCas9 alone (**Figure 3B**). VPR had no transcriptional effect, possibly because it offset the dCas9-mediated repression (data not shown). We also measured the impact of targeting this promoter in a different tissue context, to compare the results to the wing disc measurements. Because we know that Rbf2 is localized to the gonads of adults, we used the *traffic jam*-GAL4 driver to express the dCas9 chimeras in the follicle cells of the adult ovary. Using the same gRNAs, 4 + 5, we recruited Rb chimeras to the *E2F2/Mpp6* bidirectional promoter, and measured gene expression changes in whole ovaries (**Figure 3C**). Consistent with the wing disc results, the Rb-specific effects (compared to dCas9 alone) for *Mpp6* were greater than for *E2F2*. As in the wing disc, Rbf1 again was a better repressor than Rbf2 on this locus.

**Figure 3.**
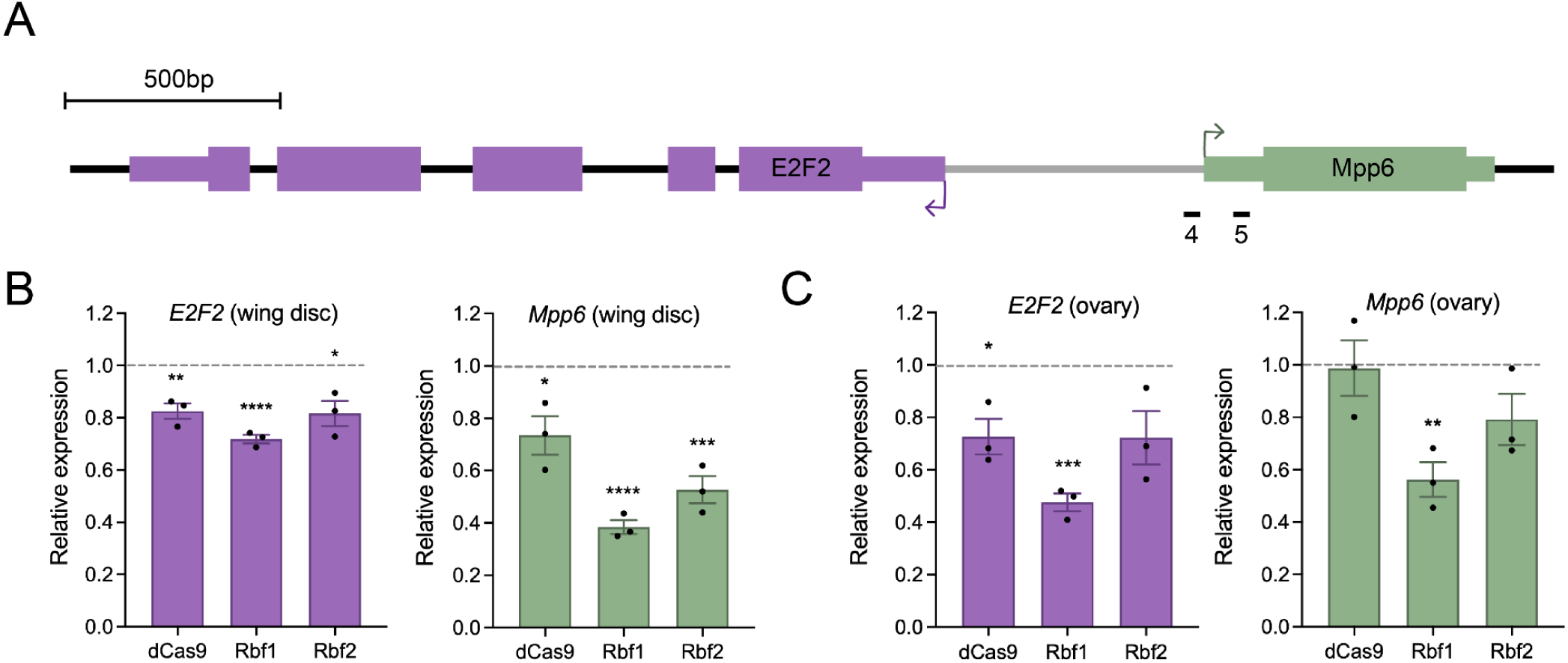
Rbf1 and Rbf2 differentially repress the *E2F2* and *Mpp6* genes in wing discs and ovaries. **A)** Schematic of the *E2F2/Mpp6* locus in the fly, with *E2F2* indicated in purple and *Mpp6* in green. The two tandem gRNAs from **Figure 2A** are indicated now as 4 and 5. **B)** L3 wing discs were dissected for RT-qPCR analysis after targeting this locus. *E2F2* and *Mpp6* expression were significantly repressed by Rb paralog recruitment. In both cases, Rbf1 represses more than Rbf2. dCas9 repressed as well, but the level of repression was not enough to create a phenotype. The control samples (set to 1) are wings in which dCas9 was targeted using the QUAS gRNA, which does not target the Drosophila genome (Ewen-Campen et al. 2017). **C)** Ovaries were collected from 3-4 day old females for RT-qPCR analysis after targeting using gRNAs 4 + 5 and the *traffic jam*-GAL4 driver. As with the wing discs, Rbf1 is a better repressor in the ovary, and both genes show significant repression. Error bars indicate SEM. * p<0.05, ** p<.01, *** p<.001, **** p<.0001.

We also measured the transcript levels of *wg*, where Rb paralogs did not show developmental phenotypes, while VPR did. As expected, *wg* levels were significantly upregulated by VPR, with 3-fold higher expression than control (**Figure S3A**). Unexpectedly, *wg* was significantly repressed by both Rbf1 and Rbf2, with Rbf1 effects slightly greater than those of Rbf2, similar to the pattern noted on *E2F2*/*Mpp6*. *wg* is a gene for which Rb ChIP peaks were not previously identified in embryos; while Rb may not be recruited to this promoter by endogenous E2F proteins, our results show that ectopic recruitment of these corepressors via dCas9 does lead to Rb-mediated gene repression (Wei *et al*. 2015). This repression seems to not disrupt wing development, while *wg* upregulation by VPR does.

We additionally measured transcript levels of *InR* after targeting its promoter. For *InR*, we did not measure a decrease in expression as we expected for dCas9 and dCas9-Rbf2, which produced adult wing phenotypes (**Figure S3B**). Instead, we measured slight activation by Rbf1 and Rbf2, while dCas9 and VPR had no effect on expression. It is possible that Rb protein recruitment on this locus interferes with an endogenous repressor. As was observed with *wg*, the transcriptional effects early in development do not correlate with the phenotypic impact in the adult stage. It is clear that in this context, there is no evidence of repression of *InR* by dCas9-Rb chimeras. Taken altogether, simply positioning an Rb protein close to a gene’s TSS does not guarantee repression; yet, we observe that in many contexts, transcriptional repression is an Rb-dependent process. What is it about the promoter that allows for Rb-mediated repression activity? To answer this question, we tested whether the site of recruitment and distance from the TSS play an important role in repression potency.

### Position-sensitive Rb paralog repression on the *E2F2/Mpp6* bidirectional promoter

*In vivo,* Rb proteins tend to bind proximal to a gene’s TSS; for Rbf1 and Rbf2, enrichment is observed at -200 bp (Acharya *et al*. 2012; Korenjak *et al*. 2012; Wei *et al*. 2015). Rb may bind in a promoter-proximal position simply because E2F proteins must work from a promoter-proximal position, or because Rb repression complex formation may require promoter proximity for optimal repression. The fact that Rb-mediated repression of *Mpp6* is greater than *E2F2* in both wing discs and in ovarian follicle cells may be a consequence of the gRNAs being positioned closer to the *Mpp6* TSS. To test whether Rb distance to the TSS affects its repression ability of these two genes we designed nine gRNAs spanning the intergenic region and UTRs. With these gRNAs, we could measure whether moving the gRNAs closer to the *E2F2* TSS would have a greater impact on *E2F2* while relieving *Mpp6* repression (**Figure 4A**). We generated new transgenic fly lines expressing one of the nine single gRNAs, and crossed them to the *nub*-GAL4>UAS:dCas9-Rb homozygous flies to generate triple transgenic flies for the experiments described below.

**Figure 4.**
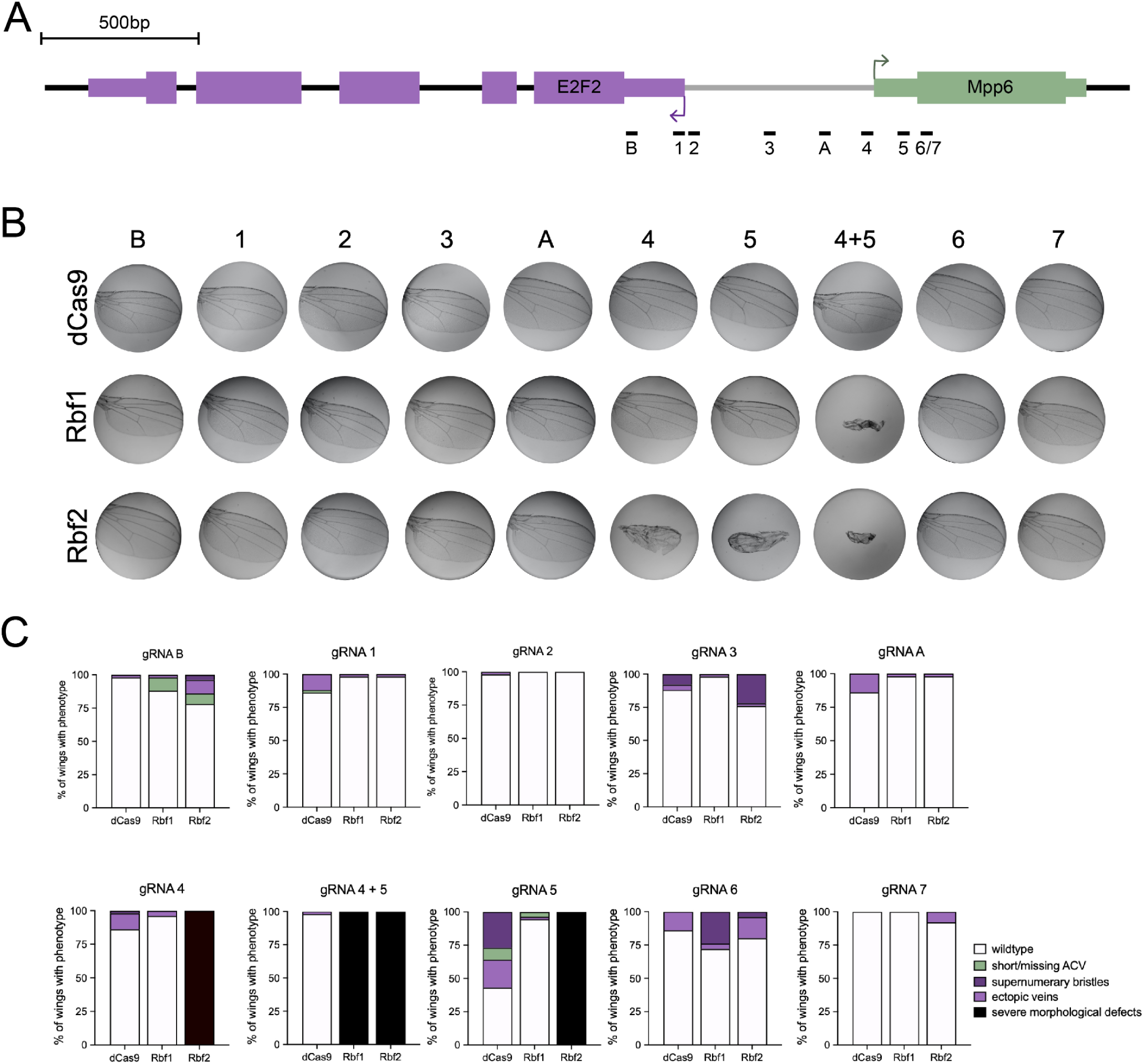
Repression of *E2F2/Mpp6* in a distance-dependent manner in the wing. Adult wings were dissected after targeting by dCas9, dCas9-Rbf1 or dCas9-Rbf2 using gRNAs designed to bind to different positions on the *E2F2/Mpp6* bidirectional promoter. **A)** Schematic of gRNA positions on the *E2F2/Mpp6* promoter used *in vivo* in the fly. dCas9 effectors were recruited to these sites using one gRNA at a time. **B)** Representative wing images from recruiting dCas9, dCas9-Rbf1 or dCas9-Rbf2 to different positions on the *E2F2/Mpp6* promoter. **C)** Graphs indicating the % of wings with particular phenotypes.

We first screened for a phenotype in the adult wings. Strong developmental effects were observed only with gRNAs in positions 4 or 5 (**Figure 4B, C**). Unexpectedly, the severe morphological defect seen with 4 + 5 in combination was only observed with the single gRNA 4 or 5 when targeting with Rbf2, but not with Rbf1. In contrast, the other gRNAs generated mild or no phenotypic effects (**Figure 4B, C**). The lack of biological activity with the single gRNAs is not simply because they are incapable of targeting this locus, as we demonstrate below with reporter assays. Additionally, recruiting the effectors closer to the *E2F2* TSS using gRNA 1 or 2 did not lead to a severe phenotype, suggesting that the severe phenotype observed with 4 + 5 may be caused by repression of *Mpp6*.

Next, we used RT-qPCR to measure changes in *E2F2* and *Mpp6* expression after targeting the effectors to each of these gRNA positions (**Figure 5**). We anticipated the greatest repression of *Mpp6* from near its TSS, with potential de-repression from other sites. Indeed, for *Mpp6*, positioning the chimeric proteins at position 4 or 5, or 4 + 5 had the greatest effect (**Figure 5C, E**). With single gRNA 4 or 5 alone, Rbf2 was more potent in repressing *Mpp6*; however, Rbf1 did cause modest repression as well. Moving the Rb paralogs further 5’ from the *Mpp6* TSS, the gene is de-repressed in a distance-dependent manner (**Figure 5C, E**). Additionally, weak, non-specific effects were observed at position 6, and no effects from position 7, ∼50 bp downstream of gRNA 5. Interestingly, from position 1, we measure statistically non-significant but apparent upregulation of *Mpp6* by both paralogs. Taken together, these results on *Mpp6* indicate that optimal repression by Rb paralogs is observed from promoter proximal sites, consistent with a short-range, and possibly promoter-specific, mechanism of repression, a functional aspect of these corepressors that was previously unappreciated. A possible advantage of such a short-range repression mechanism is that gene expression can be adjusted in a more nuanced fashion, with subtle but quantitative changes to expression.

**Figure 5.**
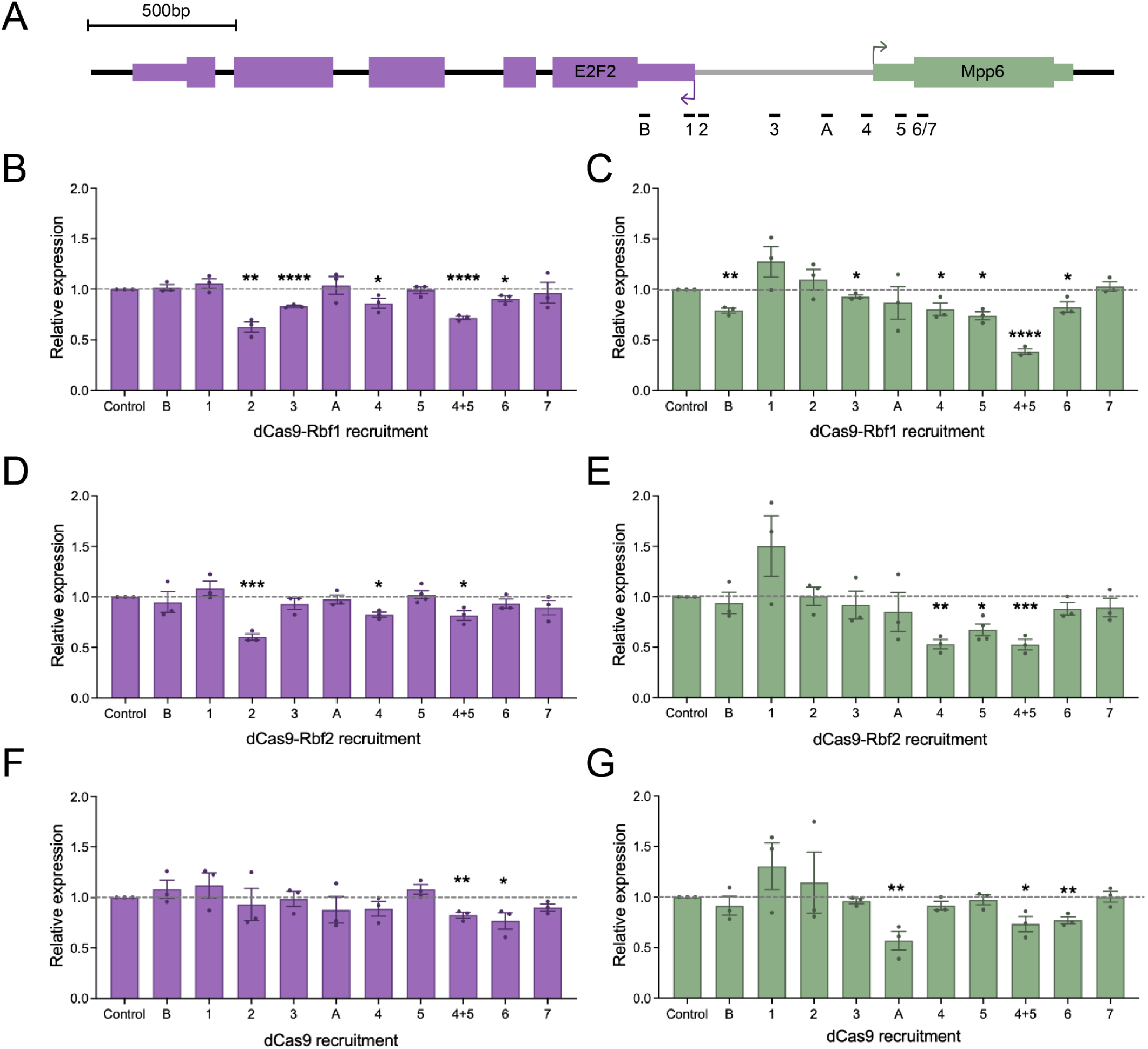
Rb regulation of *E2F2* and *Mpp6* from different positions on the promoter. **A)** Schematic of gRNA positions on the *E2F2/Mpp6* promoter used. dCas9 effectors were recruited to these sites using one gRNA at a time, and L3 wing discs were dissected for RT-qPCR analysis. **B)** Expression of *E2F2* after recruitment of dCas9-Rbf1 to each gRNA site. **C)** Expression of *Mpp6* after recruitment of dCas9-Rbf1 to each gRNA site. **D)** Expression of *E2F2* after recruitment of dCas9-Rbf2 to each gRNA site. **E)** Expression of *Mpp6* after recruitment of dCas9-Rbf2 to each gRNA site. **F)** Expression of *E2F2* after recruitment of dCas9 to each gRNA site. **G)** Expression of *Mpp6* after recruitment of dCas9 to each gRNA site. Error bars indicate SEM, and * indicates p<0.05, ** p<.01, *** p<.001, **** p<.0001.

The *E2F2* gene, while it was repressed modestly from 4 + 5, was most significantly repressed from position 2, which is also adjacent to its TSS (**Figure 5B, D**). Interestingly, the nearby position 1 did not have any effect, and neither did most other sites. Like *Mpp6*, it appears that *E2F2* is repressed by Rb paralogs through a short-range mechanism, possibly through direct contacts with transcription factors or the RNA polymerase itself, which has previously been observed *in vitro* (Ross *et al*. 1999). Few of the other sites led to *E2F2* repression, aside from the Rbf1 and dCas9-specific repression from position 6, which is puzzling. Perhaps there are elements in region 6 that regulate this gene’s expression. The fact that position 2 leads to significant repression of *E2F2* but no change in *Mpp6* expression, coupled with the lack of phenotypic effects at this site, is further evidence that the severe wing phenotype is linked to *Mpp6* repression. The results on this chromosomally embedded locus must be considered in light of the chromatin environment at this site, which is rich in TF binding and histone marks (**Figure S4**). The chromatin environment may play an important role in how this shared regulatory region is read out by transcriptional machinery and Rb proteins.

### Position-sensitive Rb paralog repression in cell culture

Initial studies of mammalian Rb relied on *in vitro* observations, and on measuring changes in gene expression after Rb perturbations. Even more recent high-throughput analysis of Drosophila Rb paralog activity has relied on uncovering differential impact on enhancers by using non-integrated reporters in S2 cell culture (Jacobs *et al*. 2022, *bioRxiv*). To compare our *in vivo* results, which are the first of their kind in the Drosophila system, to commonly used transient transfection assays in cell culture, we turned to Drosophila S2 cells. We generated two luciferase reporters using the *E2F2* and *Mpp6* promoter sequences, and recruited dCas9-Rb effectors across the promoters using the gRNAs (**Figure 6A, E**).

**Figure 6.**
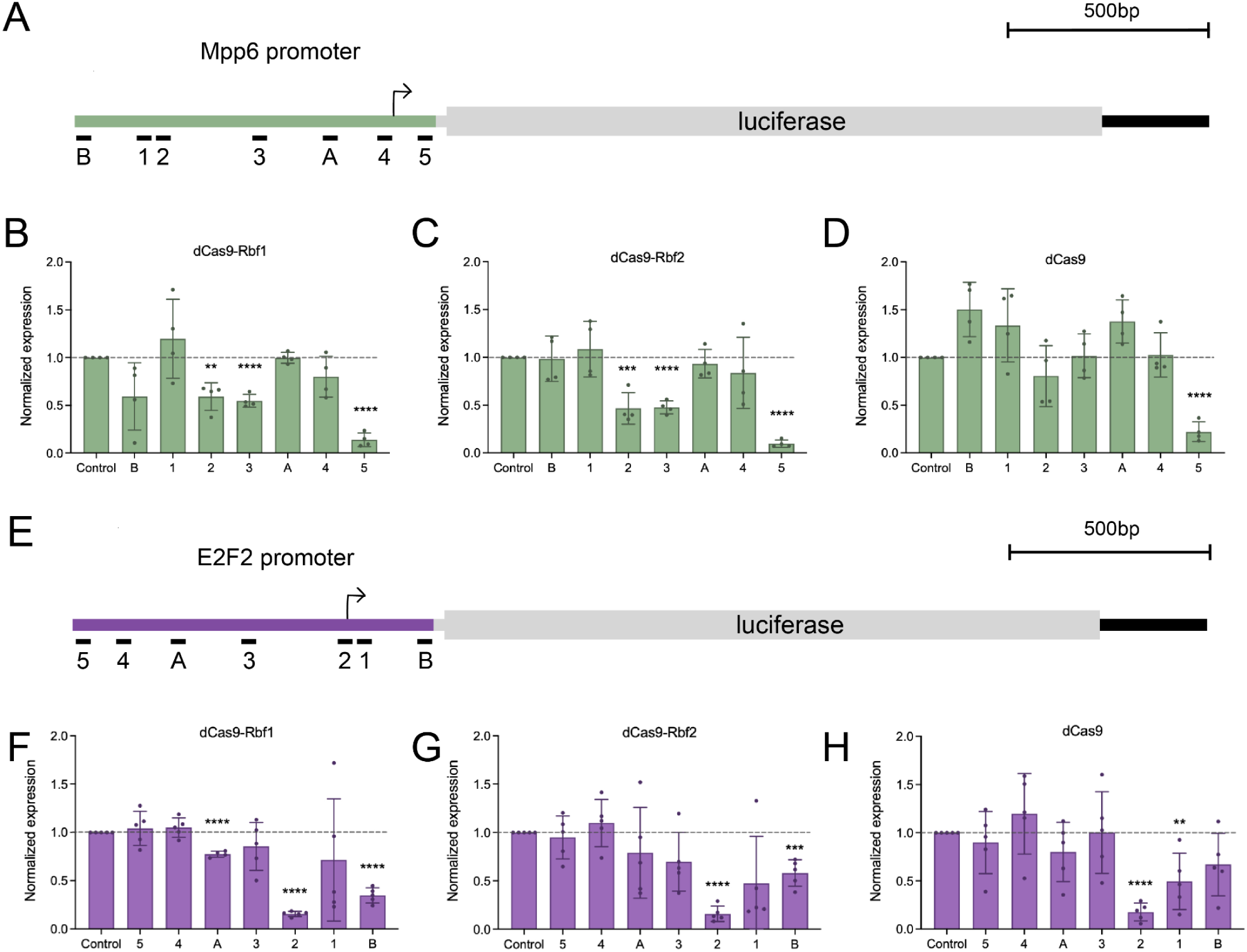
Testing positional effects of recruiting Rb paralogs to the *Mpp6* and *E2F2* promoters in S2 cells. For all experiments described here, S2 cells were transfected with *actin*-GAL4, a luciferase reporter, one of the dCas9 effectors, and a single gRNA. **A)** Schematic of luciferase reporter that was designed to be regulated by the *Mpp6* promoter. **B)** dCas9-Rbf1 has position-specific effects. Position 5 caused the same level of repression as dCas9 alone (D), suggesting steric hindrance. **C)** dCas9-Rbf2 has a similar pattern of repression as dCas9-Rbf1. **D)** dCas9 has little effect on this promoter aside from position 5, which suggests steric hindrance. **E)** Schematic of luciferase reporter that was designed to be regulated by the *E2F2* promoter. Here, the gRNAs are in opposite orientation from what is shown in panel A. **F)** dCas9-Rbf1 has position-specific effects. Position 2 caused the same level of repression as dCas9 alone (H), suggesting steric hindrance. **G)** dCas9-Rbf2, has a similar pattern of repression as dCas9-Rbf1. **H)** dCas9 has little effect on this promoter, aside from position 2, 1, and possibly B, which suggest that this promoter is more sensitive to recruitment of a large protein than the promoter in the opposite orientation. Error bars indicate SEM, and * indicates p<0.05, ** is p<.01, *** p<.001, **** p<.0001.

Given the wing disc results, we anticipated repression from positions 4 and 5 for the *Mpp6* reporter. Instead, we measured the strongest repression from positions 2 and 3, which bind at around -600 and -400 bp from the *Mpp6* TSS, respectively (**Figure 6B, C**). In this assay, Rbf1 and Rbf2 had an almost indistinguishable effect, and were both able to repress from sites distal to the TSS in this assay. We also measured significant repression from position 5, an effect that was observed with dCas9 alone as well, suggesting steric hindrance from dCas9 complexes located 3’ of the TSS (**Figure 6D**). These S2 cell data suggest that the differences in the chromatin environment between the endogenous target genes in the natural chromosomal environment and reporter genes in transiently transfected plasmids have an impact on Rb protein function, although the differences in cell types must also be considered. Additionally, the quantitative differences between Rbf1 and Rbf2 observed in the wing system are not fully recapitulated in the transfection assay, indicating that studies in a developing tissue are important for understanding the developmental impacts of Rb-mediated gene regulation, which are not measurable in transient transfections.

Interestingly, the effects observed from targeting the dCas9 proteins to the *Mpp6* reporter are distinct from those observed for the *E2F2* reporter, which is driven by the same cis-regulatory region, but in the opposite orientation (**Figure 6E-H**). The nonspecific CRISPRi effect at gRNA 5 is absent for *E2F2*, while gRNA 2, which lies close to the *E2F2* TSS, now shows non-specific interference. Not surprisingly, gRNA 1 and B, which are immediately 3’ of the *E2F2* TSS, also mediate nonspecific repression. Regulation from gRNA 3 is still Rb-specific, but weaker than for *Mpp6*. The reasons are evident for differential responses to promoter-obstructing dCas9 molecules, considering the different TSS for *Mpp6* and *E2F2*; however, the different impact of targeting site 3 suggests that the joint *cis*-regulatory region, and possible alterations after Rb recruitment, are differently read out by each gene. Taken altogether, the range of action on the *Mpp6* and *E2F2* reporters suggests that Rb proteins can function from longer distances (∼600 bp) than was optimal for the native chromosomally integrated genes, and some of the distinctions between Rbf1 and Rbf2 in the wing disc were not evident for these reporter assays. While transient reporters may not provide a complete picture of repression potential, they remain a useful tool—these experiments provided a suitable test for the gRNAs that were inactive in the disc, such as gRNA 1, showing that the gRNA was at least capable of mediating steric blocking in S2 cells.

### Effect of non-chimeric Rbf1 and Rbf2 proteins on the *E2F2/Mpp6* promoter

Given the diverse activities of dCas9-Rb chimeras on these regulatory regions, it was important to test the effect of Rb proteins interacting with native E2F binding sites as well. Thus, we expressed Rb paralogs (not fused to dCas9) in cell culture, to measure their abilities to regulate the *Mpp6* and *E2F2-*luciferase reporters. Both Rbf1 and Rbf2 significantly repressed the *Mpp6* promoter, with stronger effects by Rbf1 (**Figure S5B**). On the *E2F2* promoter, Rbf1 significantly repressed the reporter, but Rbf2 had only very mild effects (**Figure S5E**). Intriguingly, for this divergently transcribed region, the impacts of Rbf2 differ greatly, whereas Rbf1 seems to have a similar impact on both promoters.

We identified a highly conserved E2F motif in the 5’ UTR of *E2F2*, adjacent to the Rbf1 and Rbf2 ChIP peaks in the embryo (**Figure S6B, 2A**). Removal of E2F motifs in transfected reporters inhibits the ability of mammalian Rb to repress a reporter gene, underscoring the importance of E2F motifs for Rb activity (Weintraub *et al*. 1992; Lam and Watson, 1993; Dynlacht *et al*. 1994; Qin *et al*. 1995). To determine whether this particular E2F motif plays a role in the differential impacts observed by the Rb paralogs, we mutated the E2F motif in the two luciferase reporter plasmids (**Figure S5A, D, S6A, E**). Overexpression of Rb paralogs on the *Mpp6*-luciferase ΔE2F reporter led to similar levels of repression as on the WT promoter (**Figure S5C**). Both Rb paralogs still similarly repressed, but the magnitude was somewhat dampened, indicating that for *Mpp6* expression, this E2F motif is meaningful but dispensable. Another mutant, E2F^4X^ also had the same mild effect (**Figure S6A, D**). Other reports from mammals have also indicated partial redundancy of E2F motifs on promoters (Burkhart *et al*. 2010a).

On the *E2F2*-luciferase ΔE2F reporter, repression by the Rb proteins is attenuated, with no response to Rbf2 overexpression (**Figure S5F**). In this context, the E2F motif is critical for regulation by both Rb paralogs. Thus, the relative contribution of this E2F motif for the *Mpp6* and the *E2F2* promoters differs; for the two genes, the common *cis*-regulatory sequence is read out in disparate ways. Unexpectedly, the *E2F2*-luciferase E2F^4X^ mutant was not only not repressed by Rb overexpression, but showed upregulation in some instances, possibly because the mutation may have introduced other regulatory information into the gene (**Figure S6G**).

### Functional analysis of Rbf1 features necessary for repression

Previous data indicate that Rbf1 and Rbf2 can show similar or dissimilar activities depending on the promoter context. One idea that has been adduced is that differential regulation by Rbf2 is due to divergence between the paralogs’ C-terminal domains. The Rbf1 CTD differs from that of Rbf2 in that Rbf1 contains an Instability Element (IE)—an ancestral feature of this protein family that is also found in mammalian p107 and p130 (Acharya *et al*. 2010; Sengupta *et al*. 2015). Residues within the p107 IE are responsible for E2F4-specific contacts, allowing for differential binding; therefore, this IE may be chiefly important for dictating binding specificity between Rb and E2F proteins (Liban *et al*. 2016; Liban *et al*. 2017).

In the fly, the overexpression of Rb proteins in different contexts has demonstrated that the IE can impact transcriptional activity but has not differentiated between effects on promoter binding versus inherent transcription activity. An Rbf1^ΔIE^ mutant can still interact with E2F1 and E2F2, but is unable to repress cell cycle promoters such as *PCNA* (Acharya *et al*. 2010; Raj *et al*. 2012a). Overexpression phenotypes in the fly eye and wing suggests that it may be a hypomorph (Acharya *et al*. 2010; Elenbaas *et al*. 2015). Indeed, based on RNA-seq results, there is limited repression of genes after overexpression of Rbf1^ΔIE^ in wing discs, in comparison to overexpression of the WT Rbf1 protein (**Figure S7A, B**). Still, there are some contexts in which Rbf1^ΔIE^ is a good repressor: reporter assays in S2 cells showed that Rbf1^ΔIE^ is still capable of repressing *InR*, *Wts*, and *Pi3K* (Raj *et al*. 2012a). Thus, on some gene promoters, the IE is necessary for repression, but on others it is dispensable for activity.

To test the ability of Rbf1^ΔIE^ to repress when tethered to dCas9, an experiment which clearly differentiates between targeting and differential transcriptional activity, we created a fly line expressing UAS:dCas9-Rbf1^ΔIE^ and recruited it to the *E2F2/Mpp6* promoter using gRNAs 4 + 5 (**Figure 7A, C**). Intriguingly, dCas9-Rbf1^ΔIE^ caused the same severe wing phenotype as dCas9-Rbf1, and produced similar levels of repression of *E2F2* and *Mpp6* in the wing disc (**Figure 7B, D, E**). As a dCas9 chimera, the Rbf1^ΔIE^ mutant protein is as potent as WT Rbf1; thus, on this promoter, the IE is not required for repression. These results are corroborated by transient transfections in S2 cells, where dCas9-Rbf1^ΔIE^ repressed the *Mpp6*-luciferase reporter to the same extent as dCas9-Rbf1 (**Figure S8B**). Taken together, these results suggest that the IE is not required for transcriptional repression, consistent with transfection studies (Raj *et al*. 2012a); instead, it seems to be involved in recruitment to the genome via E2F. We therefore conclude that the inability of Rbf1^ΔIE^ to repress genes in our previous Rbf1 overexpression assays is due to a defect in recruiting, rather than inherent repression activity mediated by the IE.

**Figure 7.**
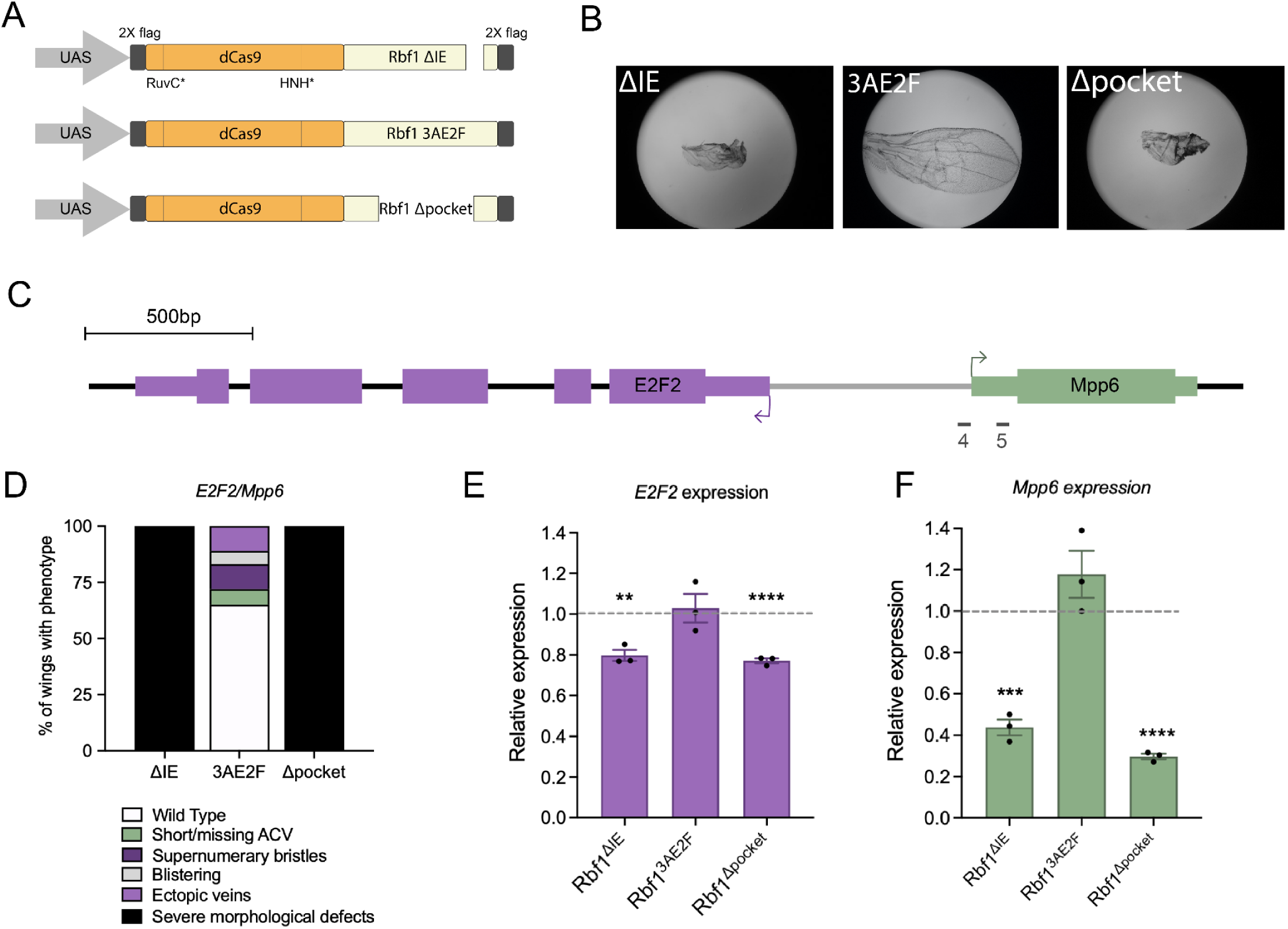
Targeting Rbf1 mutants to gene promoters indicates that the IE and pocket are not necessary for repression. **A)** Schematic of three Rbf1 mutants which were fused to dCas9 for targeting. ΔIE refers to removal of residues 728 to 786 from the C-terminus, 3AE2F is a novel E2F binding mutant, and Δpocket removes residues 376 to 728. **B)** Recruitment to the *E2F2/Mpp6* locus using the tandem gRNAs 4 + 5 led to severe morphological defects with ΔIE and Δpocket, but milder effects with 3AE2F. **C)** Schematic of the E2F2/Mpp6 promoter, with gRNAs 4 + 5 indicated. **D)** Proportion of wings with phenotypes after targeting Rbf1 mutants to position 4 + 5. **E, F)** RT-qPCR data from targeting Rbf1 mutants to gRNA positions 4 + 5. *Mpp6* expression is repressed equally well by Rbf1^ΔIE^ and Rbf1^Δpocket^, as compared to WT Rbf1 (**Figure 2B**), suggesting that the Instability Element and the pocket domain are not required for repression. However, the Rbf1^3AE2F^ mutant is unable to mediate repression, suggesting that without E2F binding, Rb cannot function as a repressor. Similar trends are seen for *E2F2*, where Rbf1^3AE2F^ is unable to repress. Error bars indicate SEM.

The C-terminal domain can influence E2F-mediated interactions; however, the pocket domain is absolutely necessary for E2F binding. The cleft between the cyclin A and cyclin B domains binds to the transactivation domain of E2F, and as discussed previously, the pocket also interacts with a variety of chromatin modifying complexes. In the fly, an Rbf1^Δpocket^ mutant only weakly represses a *PCNA*-luciferase reporter, and its interaction with E2F1 is abolished (Acharya *et al*. 2010). A plethora of studies using mammalian pocket domain mutants corroborate these findings that without the pocket, Rb cannot function as a corepressor (Bremner *et al*. 1995). Thus, it has been hypothesized that the pocket domain is key to Rb’s transcriptional repressive activities. Yet, contradicting data shows that pocket mutants that have abolished LxCxE-binding or E2F binding still function *in vivo*, putting into question the requirement of a pocket and the significance of the remaining N-and C-terminal domains (Sellers *et al*. 1998; Sun *et al*. 2006; Cecchini *et al*. 2014).

To test the significance of the pocket domain for intrinsic repression activity, we generated a fly line expressing UAS:dCas9-Rbf1^Δpocket^. Strikingly, recruitment of Rbf1^Δpocket^ to the *E2F2/Mpp6* promoter using gRNAs 4 + 5 resulted in the same severe phenotype as dCas9-Rbf1, and repressed both *E2F2* and *Mpp6* to the same magnitude as the WT protein (**Figure 7**). These results were also corroborated in cell culture, where the pocket mutant functioned similar to WT Rbf1 (**Figure S8D**). This is the first demonstration that Rb lacking the pocket domain is functional. A key takeaway from these data is that the N-and C-terminal domains, which are the more variable portions of Rb proteins, appear to be intact and functional in this context, perhaps allowing for interactions with cofactors. Indeed, the NTD resembles the cyclin folds of the pocket, and the CTD plays a role in interactions with E2F proteins; thus, they may retain the ability to create a repression complex. Interactions with cofactors have been reported outside of the pocket domain, making this a real possibility (Hassler *et al*. 2007).

We also addressed whether the Rb-E2F interactions are wholly separable from Rb interactions with cofactors by creating a mutant version of Rbf1 that lacks E2F binding but retains an intact pocket. Based on *in vitro* work testing the mammalian Rb pocket binding to E2F1, we identified three conserved residues in Rbf1 that are reported to be essential for high affinity binding to E2F proteins (Lee *et al*. 2002). Based on this work, we made three mutations (F476A, E541A, and K588A), to create an E2F-binding mutant form of Rbf1 (Rbf1^3AE2F^). We generated flies expressing UAS:dCas9-Rbf1^3AE2F^ and tested this mutant on the *E2F2/Mpp6* promoter using gRNAs 4 + 5 (**Figure 7A, C**). Targeting led to significant morphological defects in the adult wing, although not as numerous or as severe as those induced by WT Rbf1 or the other Rbf1 mutants (**Figure 7B, D**). Interestingly, this mutant protein did not show transcriptional repression of *E2F2* or *Mpp6* in the wing disc. It is possible that these particular mutations affect stability or activity of Rbf1. Yet, when tested on the *Mpp6*-luciferase reporter in S2 cells, it exhibited repression activity similar to that of WT Rbf1. Taken together, these data suggest that Rb binding to E2F does not appear to be required for repression activity, as this mutant still retains the ability to impact wing development and gene expression in cell culture (**Figure S8C**).

## DISCUSSION

### Rb paralogy

Gene duplication is associated with functional diversification among paralogs. For transcription factors and cofactors, such diversification may reflect expression changes, whereby paralogs are differentiated by promoter evolution. At the same time, there are numerous examples of changes in activity due to protein evolution, which may impact genomic targeting, or inherent transcriptional activity. Rb paralogs in vertebrates and Drosophila share common structural and functional properties, but are clearly differentiated by unique expression patterns (Jiang *et al*. 1997; Keller *et al*. 2005). Furthermore, they have unique genomic binding patterns, which may explain divergent gene regulatory effects in knock-out experiments (Chicas *et al*. 2010; Wei *et al*. 2015). The net result of the combined changes is a distinct set of regulatory outputs: p130 has a predominant regulation during G0, while p107 and Rb regulate genes in G1 phase. Depletion of each family member in human senescent cells leads to only a 5% overlap of misregulated genes, with unique sets of genes misregulated by each paralog (Chicas *et al*. 2010). Similarly, Rbf1 and Rbf2 bind to some genes in common, but many genes are occupied uniquely by one paralog, in particular Rbf2, which has about twice the number of targeted promoters (Wei *et al*. 2015). Misexpression of the paralogs accordingly produces very different gene regulatory effects. Even for the promoters bound by both Rbf1 and Rbf2, however, misexpression or knockdown of each Rb paralog usually has divergent effects, indicating that the proteins’ functions are not identical.

To what extent has protein evolution driven changes in Rb paralog activity? At the primary sequence-level, p107 and p130 share ∼50% sequence identity, but only ∼30% with Rb, with greatest divergence in the N-and C-terminal domains (Mulligan and Jacks, 1998). Divergence in the CTD is functionally significant, as it leads to differential interactions and affinity for E2F family proteins; the Rb CTD binds preferentially to E2F1, while p107 features CTD contacts that drive a preference for E2F4 (Liban *et al*. 2017). Rbf1 and Rbf2 also have differential interactions with E2F activators; Rbf1 interacts with both E2F1 and E2F2, but Rbf2 only interacts with the E2F2 repressor (Stevaux *et al*. 2002). Thus, it is possible that like the mammalian Rb proteins, the divergence in their C-terminal domains led to preferential interactions with E2Fs. By directly testing Rbf1 isoforms in a side-by-side manner with our CRISPRi approach, we demonstrated that the IE domain of the CTD is not essential for repression activity, as has been found with some transfected reporters in cell culture (Raj *et al*. 2012a). The IE likely plays a role in differential targeting - notably, the loss of the IE is a derived feature of the Rbf2 CTD.

Until now, there has been relatively little information on whether structural divergences also impact inherent transcriptional activity. For instance, Rbf1 and Rbf2 have different abilities to repress specific promoters, but this may be strictly a function of differential promoter binding (Wei *et al*. 2015; Mouawad *et al*. 2019). Some of the earliest molecular biology studies pointed to invariant mechanisms of repression. For instance, GAL4-Rb paralog chimeras act in a similar manner on reporter genes (Bremner *et al*. 1995; Ferreira *et al*. 1998; Meloni *et al*. 1999). Chimeras interchanging the Rb and p107 pockets retain function as GAL4 fusions, indicating overlapping gene-regulatory functions (Chow *et al*. 1996). Thus, most functional studies have concentrated on the similarities between the proteins. However, structural innovations among the Rb paralogs may impact actual repression potential, once bound to a locus. The recent study by Jacobs *et al*. reported an extensive comparison of corepressors, including Rbf1 and Rbf2, in their activity to modulate Drosophila enhancers in a high-throughput assay. They found that Rbf1 and Rbf2 chimeras, while not producing identical patterns of activity, were much more similar to each other than to other diverse corepressors, including CtBP, SIN3 and CoREST (Jacobs *et al*. 2022, *bioRxiv*). It is notable that our assays of dCas9-Rb repressors uncovered similarities but also important functional distinctions, particularly when assayed on endogenous targets.

The evolutionary selection that has preserved three Rb paralogs in vertebrates, and two paralogs in the Drosophila lineage, is likely to reflect effects on diverse biological processes. Does the relatively new “experiment” in Drosophila reflect a subfunctionalization, i.e. division of labor, recapitulating an ancient molecular event that occurred in our own piscine progenitors? Some lines of evidence do support a model for parallel evolution; for instance, the conserved feature of preferential binding of ribosomal protein genes by a derived Rb paralog in both flies and humans, or the loss of the IE in derived CTD of Rbf2 and (partially) in Rb itself, which may impact targeting. These specializations, in both vertebrates and Drosophila, may have allowed for certain paralogs to focus regulation on cell growth related processes, such as protein synthesis and establishment of polarity, while other paralogs predominate for an ancestral function of cell cycle control. On the other hand, Drosophila-specific phenomena may have played an important role in shaping Rb paralog structure and function; for instance, the variety of developmental cues used in diverse Drosophila to impact reproductive capacity (Sarikaya *et al*. 2019). It is notable that of the two Rb paralogs in Drosophila, Rbf2, which is highly expressed in the female ovary, and whose loss impacts reproduction, is the more evolutionary divergent of the two. More detailed studies that measure quantitative Rb paralog contributions to diverse sets of putative gene targets will build a deeper understanding of how Rb paralogy pursues novel and conserved patterns of sub-and neo-functionalization.

### CRISPRi contributions to studies of transcriptional repression

Apart from the work of the Stark laboratory, most functional analyses of Rb repression activity have involved either single gene assays or perturbation of native Rb proteins, with global assessment of the transcriptome. These latter experiments are a joint measure of protein targeting and transcriptional activity, often combining direct and indirect effects. Thus, there has been a major need for precise assessment of protein function on a greater spectrum of promoter contexts, in order to identify and characterize the context-specific effects of Rb proteins *in vivo*. An important advantage of our CRISPRi system is its ability to directly query the response of dozens of endogenous genes to Rbf1 and Rbf2 targeting, with both transcriptional as well as sensitive developmental readouts, which proved to be related, but sometimes in a complex manner. From our survey of effects at the transcriptional level, we find that most frequently, Rbf1 is a more potent repressor, although Rbf2 is more effective in specific contexts. For instance, solely Rbf2 was able to cause the severe wing phenotype when targeting *E2F2/Mpp6* using single gRNA 4 or 5, possibly because of a greater ability to work as a single nucleation site for binding of chromatin modifying factors; Rbf1 may have weaker interactions that are stimulated by positioning multiple Rbf1 molecules on the promoter. The CRISPRi approach also proved to be highly informative about the utility of transient reporter assays used in so many functional studies. By deploying the identical gRNAs and CRISPRi effectors on the *E2F2* and *Mpp6* regulatory region, we found that the reporters faithfully reflected only certain properties of regulation *in vivo*. Specific distance-dependent effects were not reproduced in this system, and a non-specific promoter-blocking action of the CRISPRi molecules was only found in transiently transfected reporters, likely because of the less defined chromatin state of these plasmid-borne genes. The transient transfection approach was a useful complement for other reasons; we were able to directly compare CRISPRi effects to regulation by free, non-dCas9-fused Rb proteins that presumably are binding E2F transcription factors. The superior repression by native Rbf1 versus Rbf2 of *E2F2* (but less so *Mpp6*) indicates that E2F targeting may influence the different response, rather than inherent transcriptional potential. Still, we argue that going forward, using CRISPRi on endogenous promoters in a living organism, as done here, will be critical for understanding physiologically relevant effects of co-repressors such as Rb.

### Soft repression by Rb

One of the striking results from this study, though perhaps disappointing from a biotechnology perspective, is that for many instances, positioning active Rb repressors near endogenous promoters had little or no effect. To a certain degree, this may reflect the robustness of the wing developmental program; transcript levels were reduced for *wg* CRISPRi, but only the dCas9-VPR activation of the gene, not its repression, impacted wing development. Out of the 28 promoters that we targeted with Rbf1 and Rbf2, only one-third (10/28) led to a wing phenotype. Even targeting *rbf1* and *rbf2* promoters showed little or no effect, although we know that the wing is impacted by perturbation in Rbf1 levels (Payankaulam *et al*. 2016). Where repression was measured, this was generally 50% or less, although that may be a reflection of the limited pouch-specific pattern of expression of the *nubbin*-GAL4 transgene in the entire wing disc. For genome engineering purposes, it may be desirable to have a repression effector, such as the KRAB domain in mammals, that uniformly and decisively silences any target. However, from the perspective of biologically-relevant regulation, such specificity and context-dependence may be exactly what is required for optimal gene regulation by these Rb corepressors.

We have suggested that Rb factors may at times act as “soft repressors” (Mitra, Raicu et al. 2021), which do not function solely as an on/off switches for gene expression, but rather modulate gene expression by dialing down expression modestly, within physiologically relevant levels. This is in line with the Stark laboratory’s experimental results that find GAL4-Rb proteins repress only half of the enhancers they are tested against, with mostly mild repression (Jacobs *et al*. 2022, *bioRxiv*). Soft repression would come into play when a target gene sits at a junction of metabolic pathways, including signaling elements such as InR and protein synthesis machinery, which are widely yet dynamically expressed in tune with demands of cellular physiology. The positional dependence and even promoter-specific effects noted for the divergent *E2F2/Mpp6* regulatory region align with a mechanism of gene regulation that is sensitive to subtleties of chromatin, neighboring factors, and basal promoter constitution.

## MATERIALS AND METHODS

### Cloning of dCas9 and gRNA constructs

The FLAG-tagged (DYKDDDDK) coding sequence for Rbf1 was obtained from the pRbf1 plasmid described previously (Acharya *et al*. 2010). Full length dCas9 (with D10A, H840A substitutions) was obtained from pcDNA-dCas9 (gift from Charles Gersbach; Addgene plasmid #47106; Perez-Pinera *et al*. 2013). The dCas9 coding sequence (with two 5’ FLAG epitope tags) was subcloned into the pRbf1 parent vector, 5’ of the Rbf1 sequence. During cloning, a linker between the 3’ end of dCas9 and 5’ end of Rbf1 was inserted, introducing PacI for simpler downstream cloning of other Rb effectors (linker sequence is 5’-GGCTTAATTAATAGTACC-3’). The dCas9-Rbf1 ORF was removed using 5’ NotI and 3’ XbaI sites and cloned into the pUASTattB vector. The final UAS:dCas9-Rbf1 vector was used to subclone other Rb FLAG-tagged sequences into the PacI-XbaI site, including Rbf2, Rbf1^ΔIE^, Rbf1^3AE2F^, and Rbf1^Δpocket^ (Acharya *et al*. 2010; Mouawad *et al*. 2020). All coding sequences for these effectors were removed from their parent vector using PCR amplification with 5’ PacI and 3’ XbaI sites introduced on either end. The dCas9 control was created by removing Rbf1 from UAS:dCas9-Rbf1 using 5’ PacI and 3’ XbaI, and inserting a gBlock containing an identical 3’ FLAG found in Rbf1 into the 3’ end of dCas9. Rbf1^3AE2F^ was created here through site directed mutagenesis (Expand Long Template PCR system) to sequentially introduce F476A, E541A, and K588A into the Rbf1 coding sequence. Single gRNAs for the *E2F2/Mpp6* promoter were designed using the DRSC Find CRISPRs tool (www.flyrnai.org/crispr/) and cloned into the pCFD3.1 vector (Addgene #123366), cutting with BbsI and inserting the annealed oligos.

### Transgenic flies

Flies were fed on standard lab food (molasses, yeast, corn meal) and kept at room temperature in the lab, under normal dark-light conditions. The *nubbin*-GAL4 fly line was obtained from the Bloomington Drosophila Stock Center (BDSC; #25754), and *traffic jam*-GAL4 was a gift from Sally Horne-Badovinac. GAL4 lines were maintained as a homozygous line with a Chr 3 balancer obtained from BDSC #3704 *(w[1118]/Dp(1; Y)y[+]; CyO/Bl[1]; TM2, e/TM6B, e, Tb[1]*). Homozygous dCas9-effector flies were generated by using the ϕC31 integrase service at Rainbow Transgenic Flies Inc. #24749 embryos were injected with each dCas9-effector construct to integrate into Chr 3 landing site 86Fb. Successful transgenic flies were selected through the mini-white selectable marker expression in-house, and maintained as a homozygous line with Chr 2 balancer (from BDSC #3704). *driver*-GAL4 and dCas9-effector flies were crossed to generate double homozygotes (*driver*-GAL4>UAS:dCas9-Rb). These flies were maintained in the lab and used for crossing to homozygous gRNA flies described below. *nub*-GAL4>UAS:dCas9-Rbf1 homozygous flies had a notched wing phenotype, consistent with Rbf1 overexpression in the wing (Elenbaas *et al*. 2015). *nub*-GAL4>UAS:dCas9-Rbf2 did not have a wing phenotype, and neither did any of the other homozygous fly lines. Two Rbf1 mutant lines (Rbf1^K774A^ and Rbf1^3SA^) were also generated but not used further in this study, as the homozygous flies had severe morphological wing defects, which did not permit us to cross them to gRNA lines (Coding sequences described in Acharya *et al*. 2010, Zhang *et al*. 2014). The dCas9-VPR construct was obtained as a fly line from the BDSC #67055 (*w[*]; P{w[+mW.hs]=GawB}nubbin-AC-62; P{y[+t7.7] w[+mC]=UAS-3xFLAG.dCas9.VPR}attP2*). This dCas9 differs from the dCas9 used in our study, as it has two additional point mutations aside from D10A and H840A. sgRNA fly lines were created by Harvard TRiP, and obtained from the BDSC (fly line numbers indicated in **Supplementary Table 1**). For gRNAs designed in this study (**Supplementary Table 2**), #25709 embryos were injected with each gRNA plasmid through the ϕC31 integrase service at Rainbow Transgenic Flies Inc., to integrate into Chr 2 landing site 25C6, which is the same landing site as the TRiP gRNA lines. Successful transgenic flies were selected through the mini-white selectable marker expression in-house, and maintained as a homozygous line. Homozygous *nubbin*-GAL4>UAS:dCas9-Rb flies were crossed to homozygous gRNA flies to generate triple heterozygotes (-/-; *nubbin*-GAL4/sgRNA; UAS:dCas9-Rb/+) that are used for all fly experiments described here.

### Genotyping flies

All flies generated in this study were genotyped at the adult stage. Flies of each genotype were homogenized (1 fly/tube) in 50 µl squish buffer (1M Tris pH 8.0, 0.5M EDTA, 5M NaCl with 1µl of 10mg/mL Proteinase K for each fly). Tubes were set at 37℃ for 30 minutes, 95℃ for 2 mins, centrifuged at 14,000 RPM in an Eppendorf Centrifuge 5430 for 7 minutes, and stored at 4℃. Following PCR amplification, amplicons were cleaned using Wizard SV-Gel and PCR Clean-Up System, and sent for Sanger sequencing.

### Imaging adult wings

Adult wings were collected from ∼50 male and female 1-3 day-old adults. They were stored in 200 proof ethanol in −20°C until mounted. Wings were removed, mounted onto ASi non-charged microscope slides using Permount, and photographed with a Canon PowerShot A95 camera mounted onto a Leica DMLB microscope. Images were all taken at the same magnification (10X) and using the same software settings.

### Wing disc dissections and RT-qPCR

50 third instar wing discs were dissected from L3 larvae and placed in 200 µl Trizol (Ambion TRIzol Reagent) and stored in −80°C until use. RNA was extracted using chloroform and the QIAGEN maXtract High Density kit, and stored in −80°C. cDNA synthesis was performed using applied biosystems High Capacity cDNA Reverse Transcription Kit. RT-qPCR was performed using SYBR green (PerfeCTa SYBR Green FastMix Low ROX by Quantabio) and measured using the QuantStudio 3 machine by applied biosystems. Three control genes were averaged (Rp49, RpS13, CG8636) for all samples with control obtained from crossing dCas9 or dCas9 effectors to a non-targeting gRNA (QUAS). Primers used are indicated in **Supplementary Table 3**. RT-qPCR was performed on 3-4 biological replicates with two technical duplicates. Student’s t-test (two tailed, p<0.05) was used to measure statistical significance. Error bars indicate SEM.

### Ovary dissections and RT-qPCR

For determining changes in gene expression in the ovary, *traffic jam*-GAL4>dCas9-Rb flies were crossed to gRNA flies. Female progeny were mated with male progeny for 5-6 days, and were fed a normal diet supplemented with yeast. 20-25 ovaries/replicate were dissected in 0.3% Triton-PBS, and were placed in 1X PBS before transferring to Trizol. RNA was extracted as described above, was immediately treated with DNase (TURBO DNA-free kit) and with RNAse inhibitor during the cDNA synthesis stage. RT-qPCR was performed as described above.

### Luciferase reporter assays

Luciferase reporter constructs were created by amplifying yw^67^ (Mani-Telang and Arnosti, 2007) genomic DNA with primers flanking the 1kb promoter that is shared between *E2F2* and *Mpp6*, including the 5’ UTR of *E2F2* and *Mpp6* but no coding sequence. The promoter was inserted into 5’ NotI and 3’ HindIII sites in the luciferase reporter plasmid described previously (Wei *et al*. 2016) in both orientations, generating *Mpp6*-luciferase (the *Mpp6* ATG is proximal to luciferase TSS) and *E2F2*-luciferase (the *E2F2* ATG is proximal to luciferase TSS) reporters (Wei *et al*. 2015). *Drosophila* S2 cells were grown in 25℃ in Schneider Drosophila medium supplemented with glutamine (Gibco) containing 10% FBS and 1% penicillin-streptomycin, and equal amounts of plasmids (250 ng of each: luciferase reporter, *actin*-GAL4 (Addgene #24344), one dCas9-Rb constructs, and one sgRNA) were co-transfected into 1.5 million cells with QIAGEN Effectene transfection reagent. Cells were harvested 72 hours later and luciferase assay was performed using the Biotium Steady-Luc HTS assay kit. Briefly, cell media was removed after centrifugation, and cells were resuspended in PBS. The lysate was used in triplicate (70 µl of lysate added to 70 µl of luciferase reagent) and quantified using the Veritas microplate luminometer (Turner Biosystems). Values were normalized to a non-targeting gRNA control (empty pCFD3 plasmid). Three technical replicates were averaged for each biological replicate. Three to five biological replicates were compared via Student’s t-test to test for significance.

### Western blot

∼15 L3 wing discs were dissected in 1X PBS and placed in 35 µl cell lysis buffer (50mM Tris, pH 8.0; 150 mM NaCl; 1% Triton X-100). Tubes were flash frozen in liquid nitrogen and stored in −80°C. Discs were homogenized and protein was extracted by freezing, thawing, and vortexing several times, followed by boiling in Laemmli buffer. 20 µl of lysate was separated on a 4-20% resolving gel (Bio-Rad Mini-PROTEAN TGX Precast Gel #456-1094), and transferred to a PVDF membrane for analysis using α-FLAG antibody (Sigma Aldrich #F3165, 1:10,000), and α-CtBP as the loading control (DNA208; Keller *et al*. 2005). Blocking with both primary and secondary antibodies was performed in 5% milk-TBST (500mM Tris-HCl, pH 7.4, 150 mM NaCl, 0.1% Tween 20). Blots were developed using HRP-conjugated GαM and GαR secondary antibody (Pierce), and imaged using SuperSignal West Pico PLUS chemiluminescent substrate.

### E2F motif search

The E2F motif, described previously (Acharya *et al*. 2012), was identified in the E2F2 gene promoter using MAST (MEME-suite v5.0.2) using p<0.0005 cut off.

### Identification of promoter landscape

We used the UCSC Genome Browser, Flybase.org, and ChIP-Atlas to identify the chromatin environment on the *E2F2/Mpp6* bidirectional promoter in both S2 cells and L3 larval discs.

## COMPETING INTEREST STATEMENT

No competing interests are declared.

## AUTHOR CONTRIBUTIONS

DNA and AMR conceived of the project, AMR developed and conducted all experiments, PC performed adult wing dissections and wing phenotype analysis. DNA and AMR analyzed the data, and wrote and edited the manuscript. All authors approved the final version of the manuscript.

## ACKNOWLEDEGEMENTS

We would like to thank Tyler Chau and Matthew Chang for their assistance analyzing transcriptomic and bioinformatic data, Megha Suresh for generating the mutant luciferase plasmids, members of the Arnosti lab for their thoughtful suggestions, and the Michigan State University RTSF and plasmidsaurus for sequencing plasmids.

## FUNDING

This work was supported by the National Institute for General Medical Sciences grant (number *R01GM124137)* to D.N.A., and the National Institute of Child Health and Development grant (number *F31HD105410)* to A.M.R.

## Notes

### Competing Interest Statement

The authors have declared no competing interest.

### Summary of Updates

This version of the manuscript has been revised to clarify some ideas and include some figures in the supplementary materials.

