## Supplementary material for "Retinoblastoma protein activity revealed by CRISPRi study of divergent Rbf1 and Rbf2 paralogs": Raicu et al. Supplementary v2.pdf

Ana-Maria Raicu[1], Patricia Castanheira[2], David N. Arnosti[2]
[1] Cell and Molecular Biology Program, Michigan State University, East Lansing, MI
[2] Department of Biochemistry and Molecular Biology, Michigan State University, East Lansing, MI

### SUPPLEMENTARY TABLES

**Supplementary Table 1** is attached as an excel sheet.

**Supplementary Table 2.** Sequences of gRNAs designed for the *E2F2/Mpp6* bidirectional promoter

| gRNA | gRNA sequence(s) |
|---|---|
| 1 | GAAAAAAATGACATAAATGG |
| 2 | GCAGTGCGCACGAAGAATAG |
| 3 | GGACAATAAACCTTAACGAA |
| 4 | TTCAGCCTAGCTAGAAAACG |
| 5 | AAAACTAGGGCGAAACCATC |
| 6 | AGTCTCGGCTTTGATTTGGA |
| 7 | TCAAAGCCGAGACTTTCGCG |
| A | AGTTGAGCTTGTTTGTCAGT |
| B | TATAGTCATCGAGTCGATTG |

**Supplementary Table 3.** RT-qPCR primers

| Gene | Primers (Forward and Reverse) |
|---|---|
| *CG8636* | F: GATCCGCTGCTAGATCCCAC<br>R: CCCTTGTACGGGCAGTTGA |
| *Rp49* | F: ATCGGTTACGGATCGAACAAGC<br>R: GTAAACGCGGTTCTGCATGAGC |
| *Rps13* | F: GGTCGTATGCACGCTCCT<br>R: CATCTGCGTTCAGTTTCAGC |
| *E2F2* | F: GACGAGGAAGTAGATATCAAGCG<br>R: TCAAAGAACCCATCCACATCG |
| *InR* | F: ACGACAACAAAACCGTTGC<br>R: TTCACGTGATCTCAATCATGC |
| *Mpp6* | F: GCTCGGTCATTCTGCTTTTG<br>R: CTCGGCTTTGATTTGGATGG |
| *wg* | F:TCGGATTCGGGTTCAAGTTC<br>R:CACTCCTGTCGCATCTCC |

### SUPPLEMENTARY FIGURES

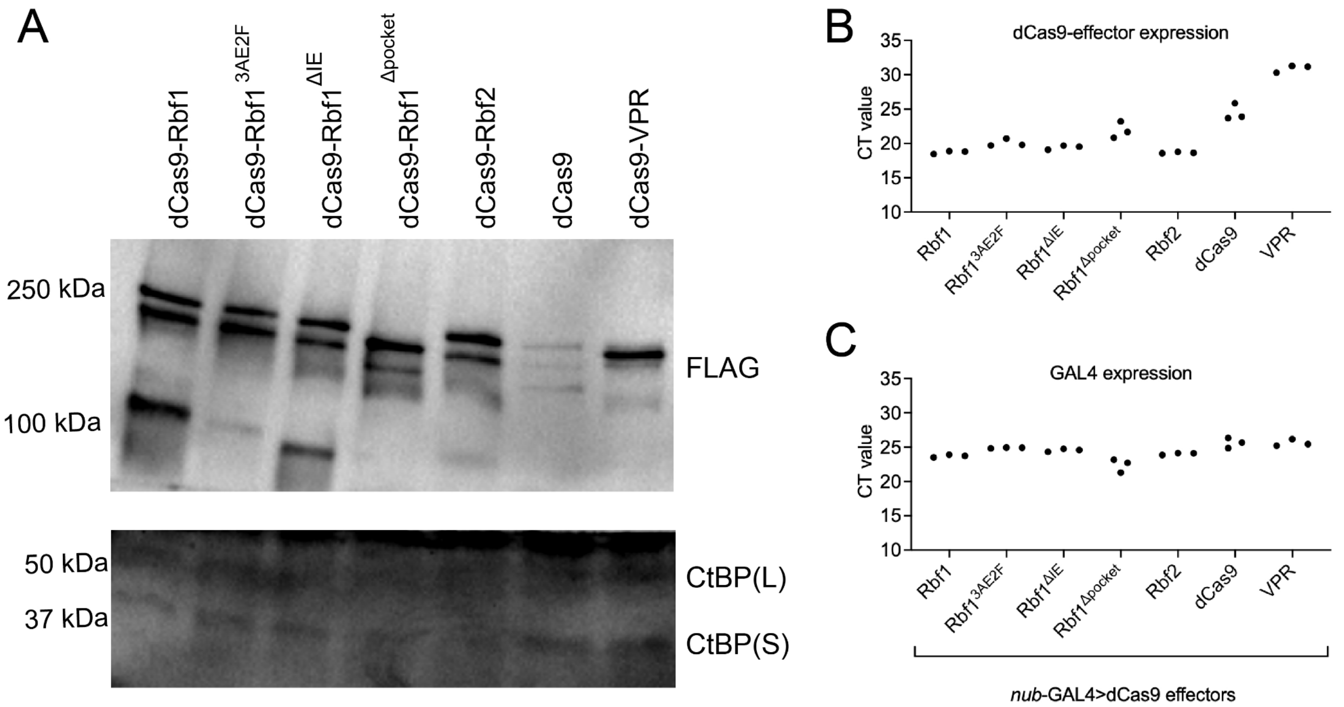


**Figure S1. Expression of dCas9 isoforms *in vivo*. A)** Western blot of dCas9 effectors expressed in L3 wing discs (genotypes shown in **Figure 1**) shows similar levels of expression of effectors at the protein level, aside from dCas9 alone, which is expressed at lower levels. Levels of endogenous CtBP(L) and CtBP(S) measured as loading controls. dCas9 proteins detected with anti-FLAG antibody, CtBP with anti-CtBP serum. **B)** mRNA levels of dCas9-Rb effectors expressed in L3 wing discs were measured using RT-qPCR. Expression level (CT value) indicates that the effectors are expressed at relatively similar levels, aside from dCas9 and dCas9-VPR which are expressed at lower levels. **C)** mRNA levels of the *nubbin*-GAL4 transgene from L3 wing discs indicates that the GAL4 driver levels are similar from one genotype to the next, as expected.

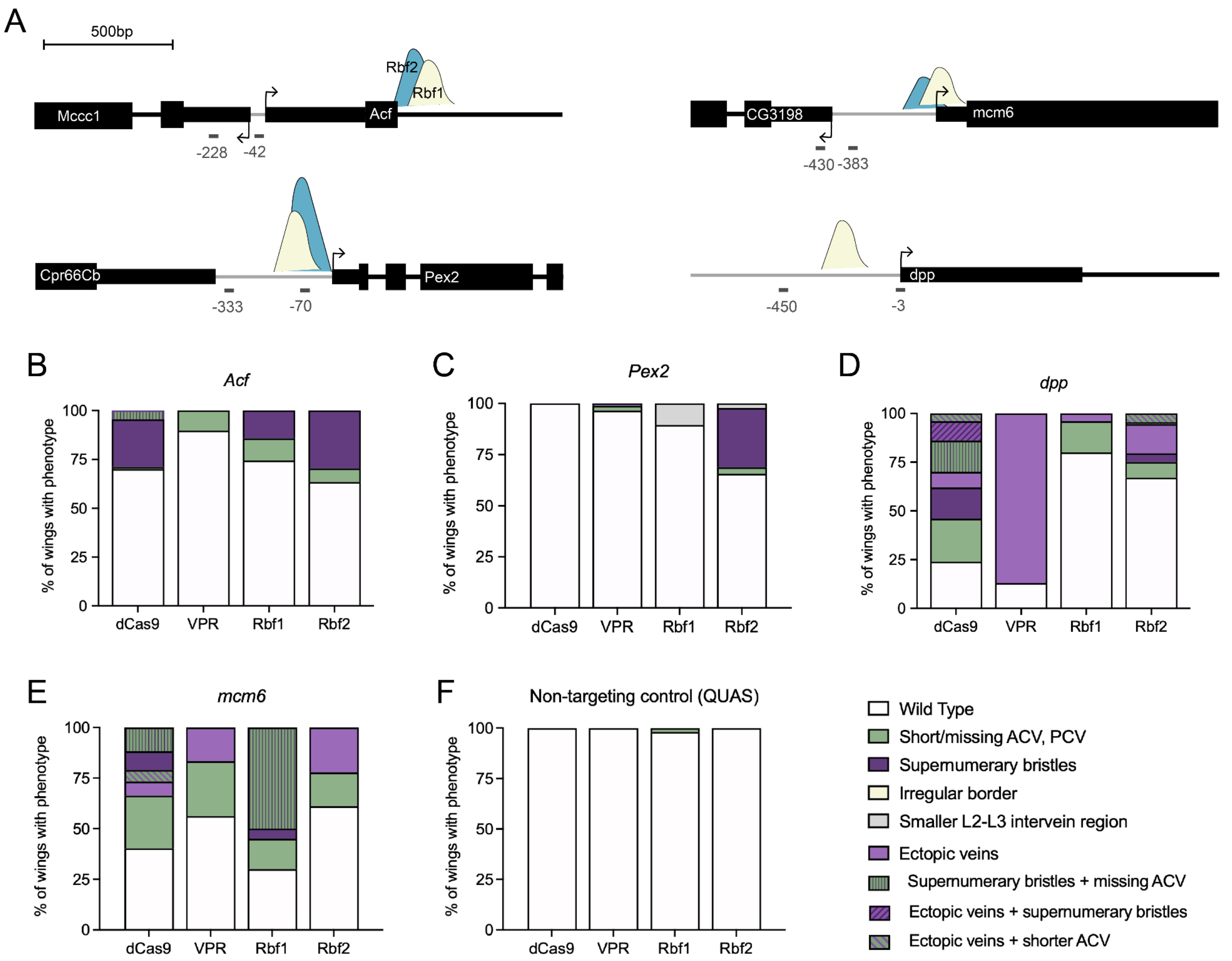


**Figure S2. Rbf1 and Rbf2 targeted to diverse gene promoters produce gene-specific effects. A)** Diagram of gRNA binding sites on the promoters of *Acf*, *Pex2, mcm6,* and *dpp*. Arrows indicate the TSS. Thick black bars are exons, with thinner regions representing UTR, and black lines are introns. Gray horizontal lines indicate intergenic regions, and gray lines below the genes are the locations of the gRNAs, with approximate distance from the TSS indicated below. These gRNAs were obtained from Harvard TRiP. Yellow peaks indicate Rbf1 binding sites and blue peaks indicate Rbf2 binding sites from embryo exo-ChIP (Wei *et al.* 2015). **B)** Targeting *Acf* caused mild phenotypes such as supernumerary bristle formation by dCas9, Rbf1, and Rbf2. **C)** Targeting *Pex2* caused mild phenotypes that are different between Rbf1 and Rbf2, but not very penetrant. **D)** Targeting *dpp* caused some effector-specific phenotypes, such as ectopic venation in almost all wings by the VPR activator. Rb paralogs had mild effects in ~25% of wings. **E)** Targeting *mcm6* caused a wide variety of phenotypes by all effectors, suggesting dCas9-induced changes, regardless of effector. **F)** Expression of effectors with a non-targeting gRNA control (QUAS) was used to determine the background of expressing dCas9-Rb in the L3 wing discs. The QUAS control demonstrates that effectors not targeted to any locus on the genome do not cause adult wing phenotypes.

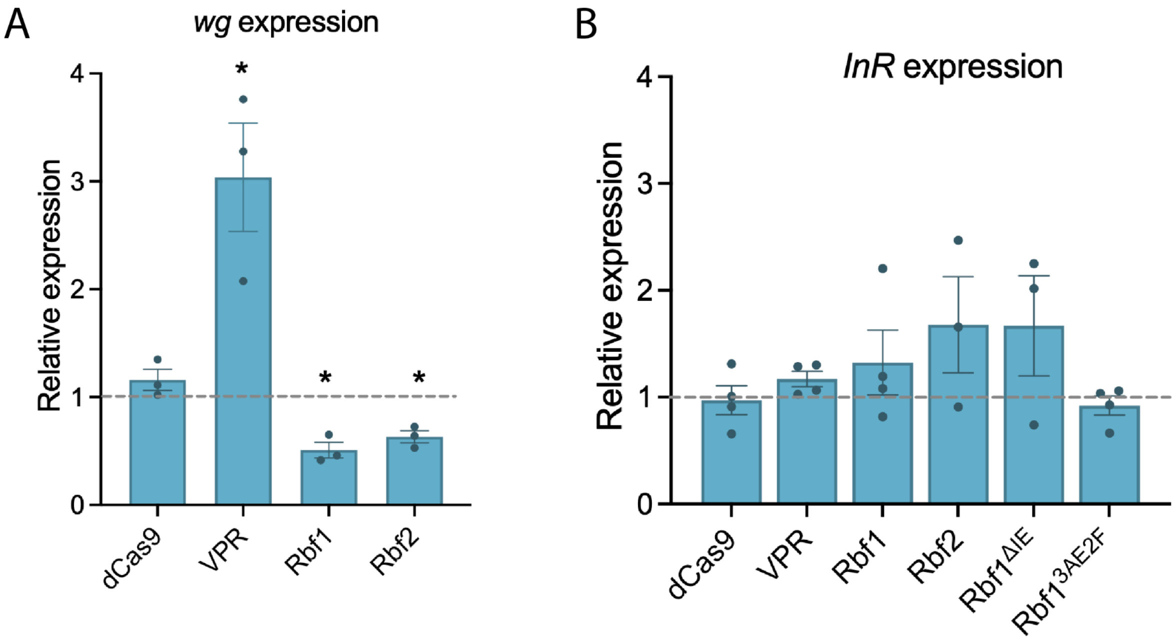


**Figure S3. Rb impact on expression of other gene targets. A)** The *wg* promoter was targeted by the Rb paralogs and the VPR activator. The VPR activator led to severe morphological defects (see **Figure 1C**), which is corroborated by a 3-4X increase in expression at the RNA level. Rbf1 and Rbf2 targeting did not lead to any overt wing phenotype, but transcript levels decreased by half. This suggests that Rb proteins can regulate at the transcript level, but depending on the gene that is being targeted, it may or may not lead to developmental defects. **B)** The *InR* promoter was targeted by Rb paralogs, Rb mutants, and the VPR activator. None of the effectors caused a decrease in *InR* expression; instead, Rbf1 and Rbf2 may have caused de-repression of the gene. $Rbf1^{\Delta IE}$ functions like the WT Rbf1, and $Rbf1^{3AE2F}$ has no effect, similar to results described in **Figure 8**. Error bars indicate SEM, and * indicates $p<0.05$.

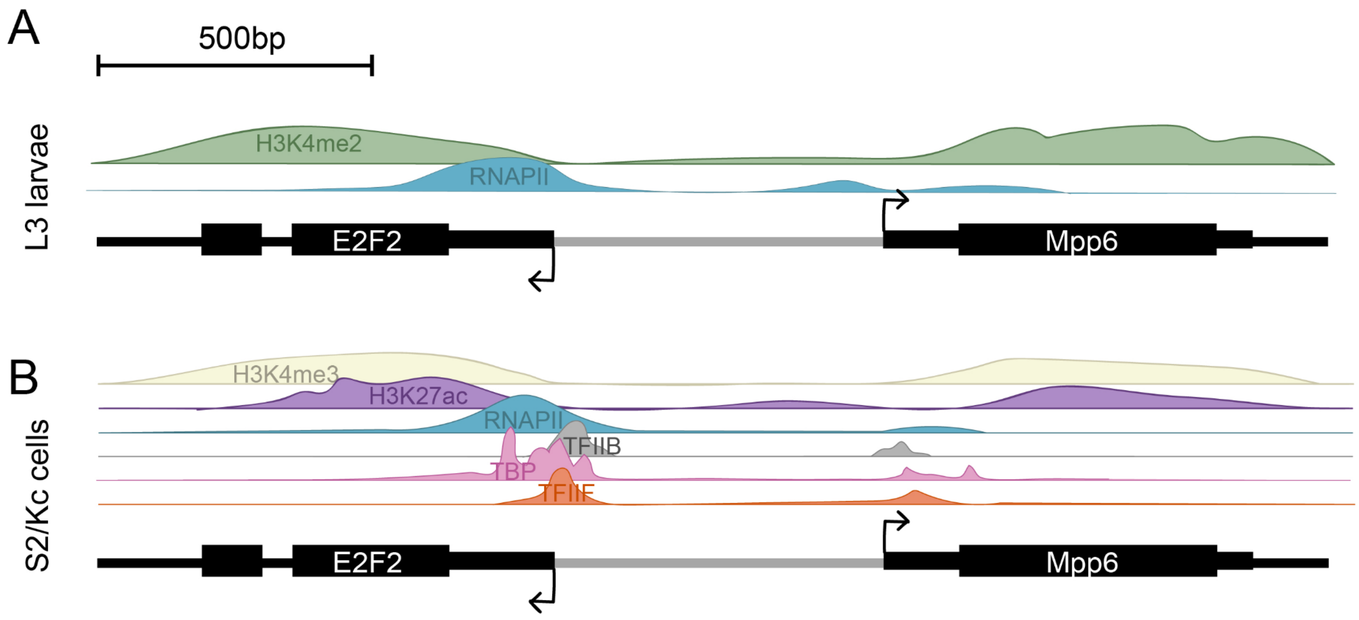


**Figure S4. *E2F2/Mpp6* promoter landscape.** Additional complexity on this locus stems from the fact that both *E2F2* and *Mpp6* are annotated to have two TSSs, each of which is about 400bp 5' of the indicated TSS (arrow). RAMPAGE-seq and CAGE-seq data indicate that the downstream (proximal) start sites are most often used, both in S2 cells and *in vivo* (UCSC Genome Browser). **A)** Chromatin landscape in L3 larvae. RNAPII binding is indicated in blue, over the two TSSs. H3K4me2 marks, which are indicative of transcriptionally active genes, are indicated in green. **B)** Chromatin landscape in S2 and Kc cells. H3K4me3 marks (yellow) and H3K27ac marks (purple) overlap the gene bodies, and are marks of active genes. RNAPII binding (blue) overlaps both TSSs, with a bigger peak over *E2F2*, where TFIIB (gray), TBP (pink), and TFIIF (orange) are also found.

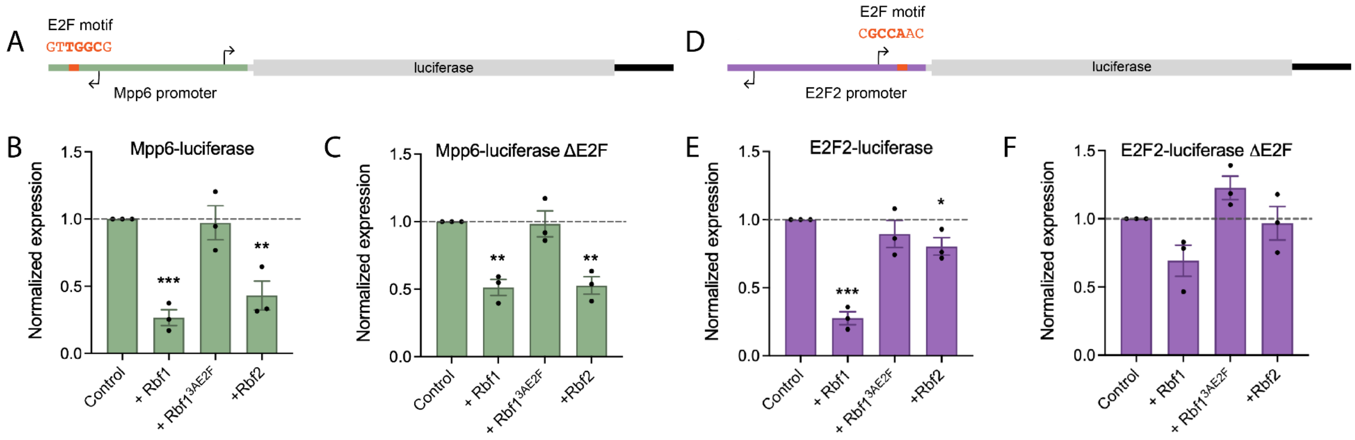


**Figure S5. Differential impact of non-chimeric Rbf1 and Rbf2 on the *E2F2/Mpp6* promoter. A)** Schematic of the *Mpp6*-luciferase reporter used in transfections with Rb plasmids. The orange box indicates a highly conserved E2F motif (**Figure S6**), with mutated nucleotides bolded in orange**.** Arrows indicate TSS. The arrow in the opposite direction indicates the *E2F2* TSS. **B)** Co-transfection of the *Mpp6*-luciferase reporter with Rbf1 and Rbf2 (untethered to dCas9) leads to significant repression by both Rbf1 and Rbf2, but not by the Rbf1$^{3AE2F}$ mutant discussed in **Figure 7**. **C)** Removal of the E2F motif from the *Mpp6* promoter (ΔE2F) does not significantly impact repression ability by Rbf1 and Rbf2, although the magnitude of repression by Rbf1 is slightly dampened. **D)** Schematic of the *E2F2*-luciferase reporter used in transfections with Rb plasmids. The orange box indicates the same highly conserved E2F motif indicated in panel A, with mutated nucleotides bolded in orange. Arrows indicate TSS. The arrow in the opposite direction indicates the *Mpp6* TSS. **E)** Co-transfection of the *E2F2*-luciferase reporter with Rbf1 and Rbf2 (untethered to dCas9) leads to significant repression by Rbf1, but not much by Rbf2, in contrast to what was seen with the promoter in the opposite orientation in panel B. **F)** Removal of the E2F motif from the *E2F2* promoter (ΔE2F) impact repression ability by both Rbf1 and Rbf2, as they are unable to significantly repress this promoter without the E2F motif intact. Error bars indicate SEM, and * indicates $p<0.05$, ** is $p<.01$, *** $p<.001$.

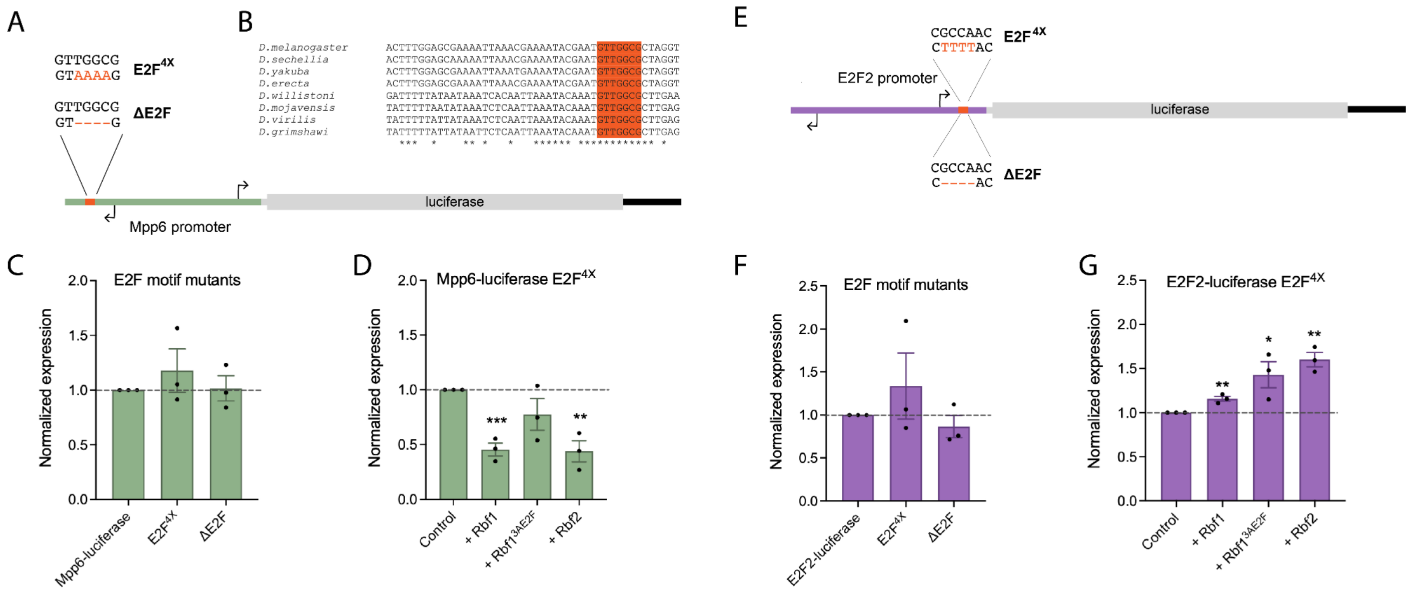

**Figure S6. The conserved E2F motif is not necessary for *E2F2* and *Mpp6* expression. A)** Schematic of the *Mpp6*-luciferase reporter used in transfections with Rb plasmids. The orange box indicates a highly conserved E2F motif that was mutated in two ways: $E2F^{4X}$ changes 4 nucleotides to A, and ΔE2F removes 4 nucleotides from the motif. **B)** The E2F motif is perfectly conserved across multiple Drosophila species. **C)** Expression levels of the two mutant promoters relative to the WT *Mpp6* promoter. Both types of mutations to the E2F motif do not affect the normal expression level of the promoter. **D)** The $E2F^{4X}$ mutant has minimal impact on how Rb proteins regulate the *Mpp6* promoter. **E)** Schematic of the *E2F2*-luciferase reporter used in transfections with Rb plasmids. The orange box indicates the same highly conserved E2F motif shown in panel B, showing the sequence of the complementary strand. The motif was mutated in two ways: $E2F^{4X}$ changes 4 nucleotides to T, and ΔE2F removes 4 nucleotides from the motif. **F)** Expression levels of the two mutant promoters relative to the WT *E2F2* promoter. Both types of mutations to the E2F motif do not appear to affect the normal expression level of the promoter. **G)** The $E2F^{4X}$ mutant abolishes Rbf1 and Rbf2's ability to repress this promoter. Error bars indicate SEM, and * indicates $p<0.05$, ** is $p<.01$, *** $p<.001$.

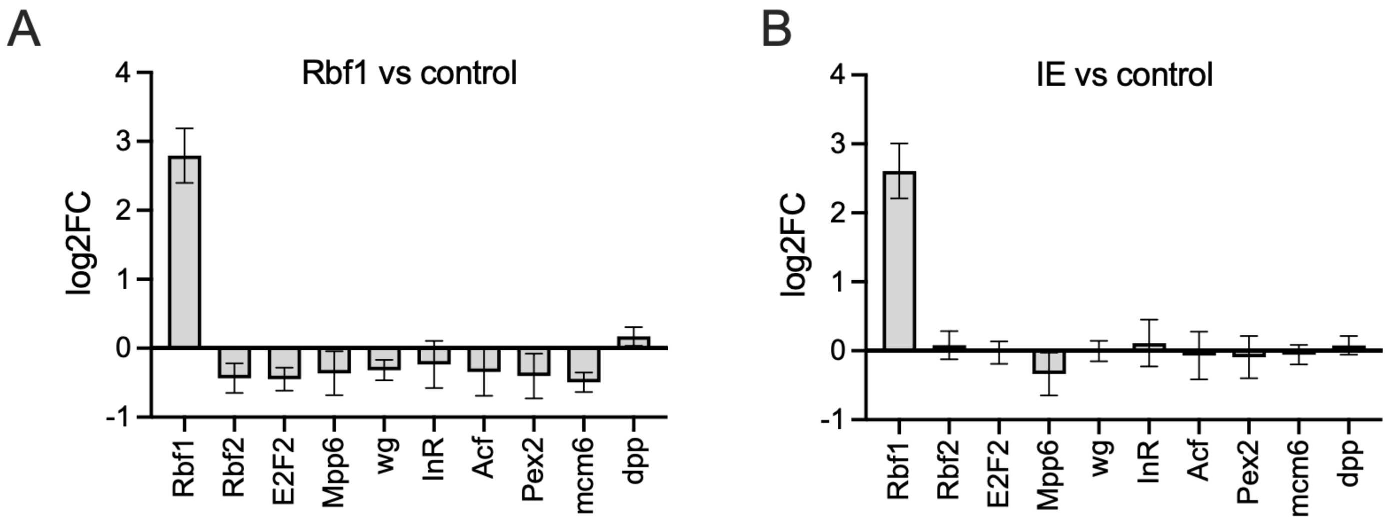


**Figure S7. Effect of removing the Rbf1 Instability Element (IE) on ability to repress gene expression in wing discs. A)** Overexpression of UAS:Rbf1 in the L3 wing disc using *pen*-GAL4 leads to modest repression of listed genes compared to the control (overexpression of UAS:GFP). **B)** Overexpression of UAS:Rbf1$^{\Delta IE}$ in the L3 wing disc using *pen*-GAL4 leads to little impact on the listed genes. These data are obtained from an RNA-seq experiment performed in L3 wing discs (See Materials & Methods in Elenbaas *et al.* 2015; Mouawad *et al.* unpublished). Rbf1 level is also shown, to illustrate that it is indeed overexpressed in this tissue.

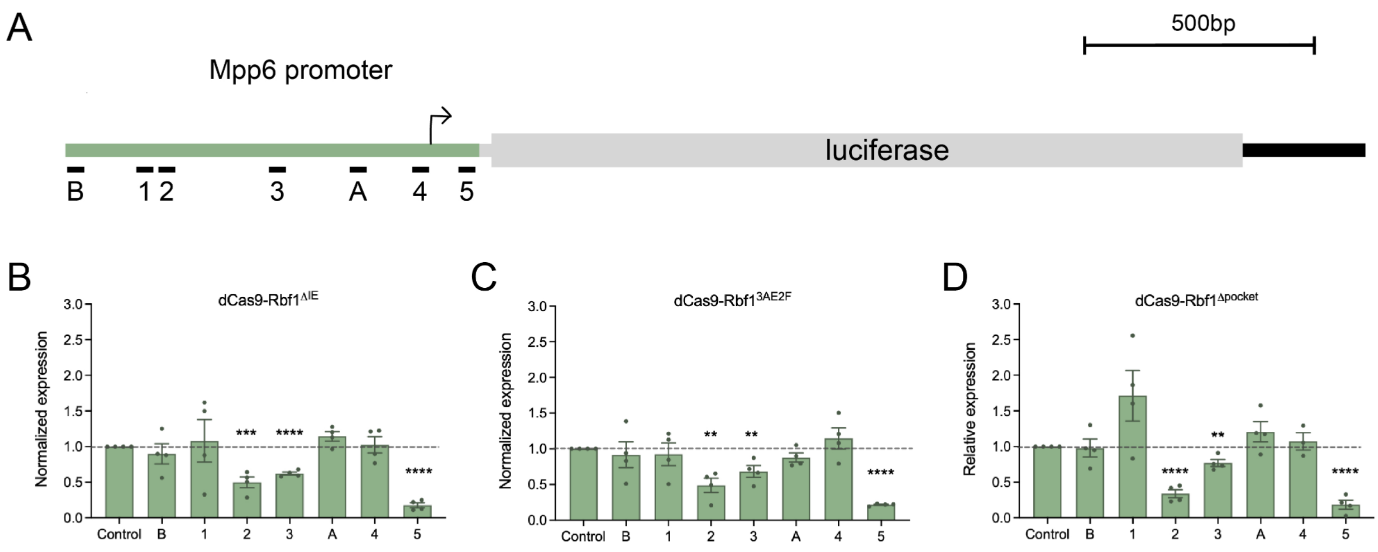


**Figure S8. Testing Rbf1 mutants in a reporter assay in S2 cells.** For all experiments described here, S2 cells were transfected with *actin*-GAL4, a luciferase reporter, one of the dCas9 effectors, and a single gRNA. **A)** Schematic of luciferase reporter that was designed to be regulated by the *Mpp6* promoter. Numbers and letters on the bottom indicate gRNA positions. **B)** dCas9-Rbf1$^{\Delta IE}$ has position-specific effects that are similar to dCas9-Rbf1 (**Figure 6**). **C)** dCas9-Rbf1$^{3AE2F}$ has position-specific effects that are similar to dCas9-Rbf1. **D)** dCas9-Rbf1$^{\Delta pocket}$ has position-specific effects that are similar to dCas9-Rbf1. Error bars indicate SEM.
